# Structural and biochemical studies define Nudt12 as a new class of RNA deNADding enzyme in mammalian cells

**DOI:** 10.1101/474478

**Authors:** Ewa Grudzien-Nogalska, Yixuan Wu, Xinfu Jiao, Huijuan Cui, Ronald P. Hart, Liang Tong, Megerditch Kiledjian

**Affiliations:** Department Cell Biology and Neuroscience, Rutgers University, Piscataway, NJ 08854; Department Biological Sciences, Columbia University, New York, NY 10027

## Abstract

We recently demonstrated mammalian cells harbor NAD-capped mRNAs that are hydrolyzed by the DXO deNADding enzyme. Here we report the Nudix protein Nudt12 is a second mammalian deNADding enzyme structurally and mechanistically distinct from DXO and targets different RNAs. Crystal structure of mouse Nudt12 in complex with the deNADding product AMP and three Mg^2+^ ions at 1.6 Å resolution provides exquisite insights into the molecular basis of the deNADding activity within the NAD pyrophosphate. Disruption of the *Nudt12* gene stabilizes transfected NAD-capped RNA in cells and its endogenous NAD-capped mRNA targets are enriched in those encoding proteins involved in cellular energetics. Furthermore, exposure of cells to metabolic stress manifests changes in NAD-capped RNA levels indicating an association between NAD-capped RNAs and cellular metabolism. Lastly, we show that the bacterial RppH protein also possesses deNADding activity toward NAD-capped RNA but not free NAD, revealing a third class of deNADding enzymes.

---

The redox cofactor nicotinamide adenine dinucleotide (NAD) was recently reported to be covalently linked to the 5’ end of bacterial RNA^1^ primarily on a subset of small regulatory RNAs in a cap-like manner^2^. Addition of the NAD cap to RNAs appears to occur during transcription initiation^3^. The transcriptional incorporation of NAD at the 5’ end is supported by *in vitro* incorporation of NAD as a “non-canonical initiating nucleotide” (NCIN) by bacterial RNA polymerase that can integrate NAD as the first transcribed nucleotide in place of ATP^3–5^. More recently, NAD-capped RNAs were also confirmed in eukaryotes with their identification in *Saccharomyces cerevisiae*^6^ and mammalian cells^7^. These findings demonstrate the broad distribution of NAD caps in diverse organisms.

The bacterial Nudix (nucleoside diphosphate linked to another moiety X) hydrolase NudC, initially described as a NAD/H pyrophosphohydrolase^8^, can remove the NAD cap by hydrolyzing the diphosphate linkage to produce nicotinamide mononucleotide (NMN) and 5′-monophosphate RNA^2^. Bacterial strains lacking NudC exhibit an increase in steady state levels of NAD-capped RNAs, interpreted to function as a stability element in bacteria^2,3^ implicating NudC as a deNADding enzyme that removes the protective NAD cap to promote decay. The NAD cap has been reported to protect RNA from 5′ end degradation by the bacterial RppH Nudix protein and RNaseE nucleases *in vitro*^2^. RppH contributes to the generation of monophosphorylated 5′ end transcripts^9^ that can further be subjected to RNaseE directed decay^10^. The mammalian Nudix protein Nudt12 is a close homolog of bacterial NudC^11,12^. It was identified as an NAD/H diphosphatase with a substrate preference for the reduced nicotinamide dinucleotide^13^. Although Nudt12 was initially shown to localize to peroxisomes when fused to a C-terminal green fluorescent protein^13^, immunocytochemistry reveals it is also cytoplasmic in at least kidney cells^14^ indicating a cytoplasmic function.

The mammalian non-canonical decapping enzyme, DXO^15^ possesses deNADding activity by removing the entire NAD moiety from the 5’ end of an NAD-capped RNA in cells^7^. Importantly, in contrast to the canonical N7 methylguanosine (m^7^G) cap, the NAD cap promotes, rather than prevents, RNA decay in mammalian cells^7^. Interestingly, removal of DXO from mammalian cells allowed detection of NAD-capped intronic small nucleolar RNAs (snoRNAs), suggesting that NAD caps can also be added to 5′ processed termini, in addition to transcriptional incorporation at the 5′ end, implying the existence of NAD capping mechanism^7,16^.

In this report, we provide several lines of evidence demonstrating that, in addition to DXO, Nudt12 is a deNADding enzyme that targets RNAs distinct from those subjected to DXO deNADding. We further demonstrate that both enzymes are functional in cells and their removal from cells significantly increased the level of NAD-capped RNA. Moreover, NAD capping can be altered by cellular exposure to stress indicating NAD cap levels can by dynamic and vary with cellular metabolic load.

## RESULTS

### Mammalian Nudt12, but not Nudt13, deNADs NAD-capped RNA *in vitro*

Eukaryotic cells possess multiple decapping enzymes which function on distinct mRNA subsets^17^. We reasoned that, analogous to decapping proteins, the recently described DXO/Rai1 family of deNADding enzymes^7,18^ may also be just one of multiple deNADding enzymes that each regulate a subset of RNAs. Two human Nudix proteins, Nudt12 and Nudt13, with reported/predicted hydrolysis activity on free NAD ^13^ were initially pursued.

To assess the potential deNADding activity of mouse Nudt12 and Nudt13, both recombinant proteins were incubated either with ^32^P labeled NAD (N_ic_pp*A; asterisk denotes ^32^P label, Fig. 1a) or *in vitro* transcribed ^32^P labeled RNA capped with NAD (N_ic_pp*A-RNA, Fig. 1b). As expected^13^, both proteins can hydrolyze free NAD into nicotinamide monophosphate (NMN; N_ic_p) and adenosine monophosphate (p*A), which can be detected by thin-layer chromatography (TLC) (Fig. 1a). Since NAD-capped RNA deNADding by Nudt12 or Nudt13 is expected to proceed through cleavage within the diphosphate of N_ic_pp*A-RNA, the released unlabeled nicotinamide mononucleotide (N_ic_p) would not be detected by TLC and the labeled p*A-RNA would remain at the origin. Therefore, the reaction products were further treated with nuclease P1, which cleaves all phosphodiester bonds within an RNA and releases the terminal labeled p*A (Fig. 1b). Interestingly, Nudt12 but not Nudt13 possessed NAD cap deNADding activity *in vitro* (Fig. 1b), demonstrating not all NAD hydrolyzing enzymes function in RNA deNADding. Moreover, Nudt12 cleavage of the NAD-capped RNA substrate between the diphosphate linkage was similar to the activity observed with NudC^2^ and distinct from that of DXO^7^. A comparison of Nudt12 decapping activity on m^7^G-capped RNA^19^ to that of its deNADding activity revealed a greater level of deNADding relative to decapping (Fig. 1c). Nudt12 deNADding activity was compromised by a catalytically inactive mutant (Fig. 1c) harboring glutamine substitutions for two glutamic acid metal coordination residues that abolish Nudt12 decapping activity^19^, demonstrating that the decapping and deNADding activities of Nudt12 utilize the same active site.

**Figure 1.**
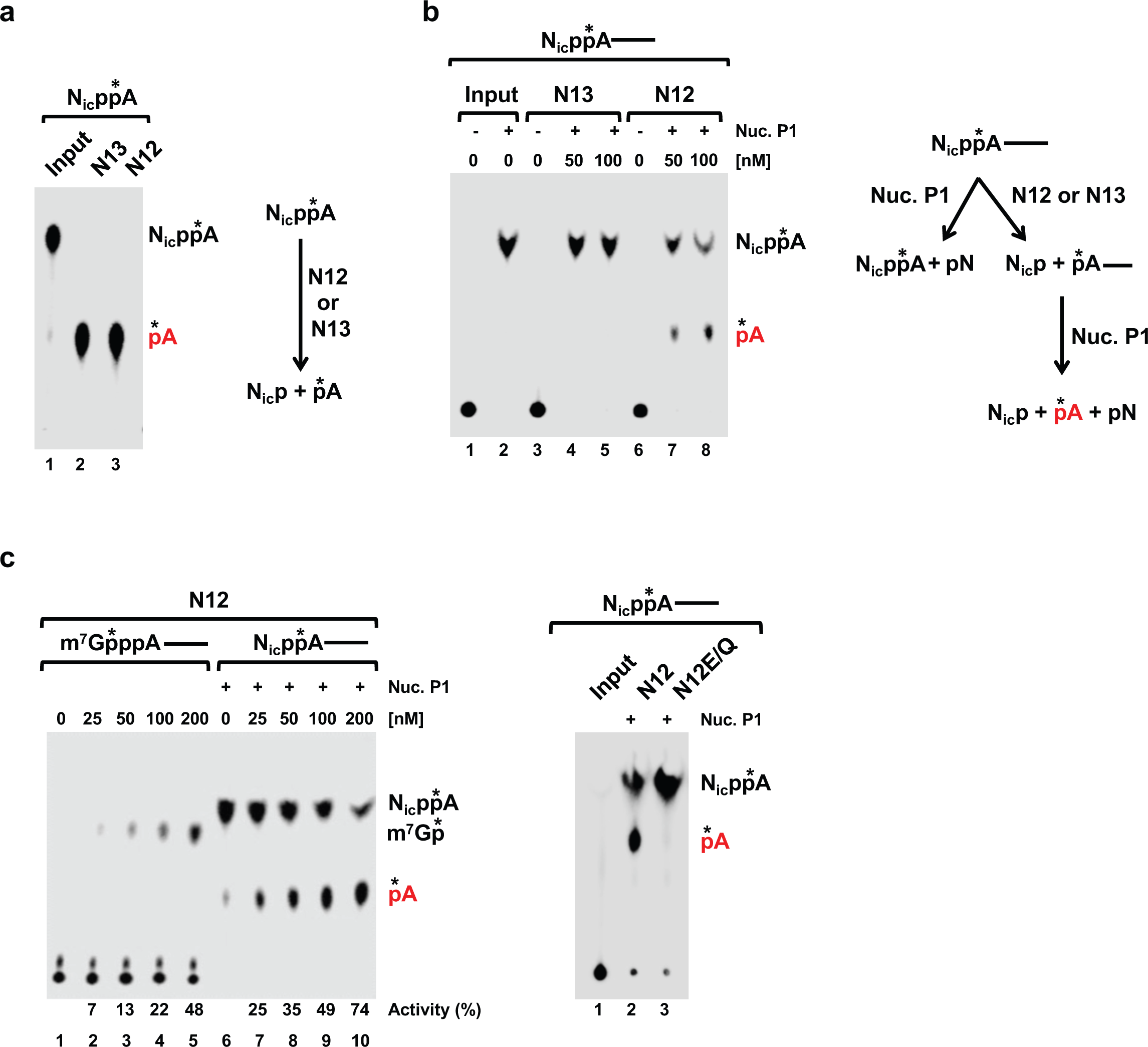
Mammalian Nudt12 possesses robust deNADding activity *in vitro*. **(a)** Mouse Nudt12 and Nudt13 can hydrolysis free NAD. Schematic of free NAD hydrolysis by Nudt12 or Nudt13 is shown on the left. N_ic_ denotes nicotinamide and the asterisk denotes position of the ^32^P. Reaction products of ^32^P-labeled free NAD (N_ic_pp*A) were resolved by polyethyleneimine (PEI)-cellulose thin-layer chromatography (TLC) developed in 0.5 M LiCl. **(b)** Nudt12, but not Nudt13 deNADs NAD-capped RNA. Schematic of NAD-capped RNA (N_ic_pp*A------) where the line represents RNA and addition of Nuclease P1 to cleave all phosphodiester bonds is denoted by Nuc.P1. Reaction products of NAD-capped RNA were resolved as in panel **a**, and (+) indicates additional treatment with Nuclease P1. **(c)** *In vitro* decapping/deNADding assays with mouse Nudt12 protein and indicated ^32^P-cap-labeled substrates. Catalytically inactive double-mutant Nudt12E373Q/E374Q (Nudt12E/Q) lacks deNADding activity.

### Distinct classes of deNADding enzymes

To determine the relative activities of Nudt12 and NudC, their hydrolysis activity on NAD, NAD-capped RNA and m^7^G-capped RNA were tested. Under these *in vitro* parameters, both enzymes possessed activity on free NAD, NAD-capped RNA and RNA capped with m^7^G (Fig. 2a). The bacterial RppH protein, which was included as a negative control, lacked hydrolytic activity on free NAD as expected (Fig. 2a), but surprisingly contained robust deNADding activity on NAD-capped RNA (Fig. 2a, middle panel) contrary to a previous report^2^. RppH deNADding activity was compromised with a catalytic inactive^19,20^ RppH mutant (Fig. 2b) demonstrating the observed deNADding activity is an intrinsic function of wild type RppH. Collectively, these findings indicate there are at least three classes of deNADding enzymes (Table 1). The first is the DXO family of proteins that remove the entire NAD from the 5’ end of an RNA^7^. The second is represented by NudC and Nudt12 which cleaves within the pyrophosphate of both NAD and NAD-capped RNA. The third class contains RppH, which does not cleave NAD, but does cleave NAD-capped RNA. The result with RppH is reminiscent of canonical Nudix m^7^G decapping enzymes that need to bind the RNA body in order to cleave the cap and contain minimal activity on cap structure alone ^19,21^.

**Figure 2.**
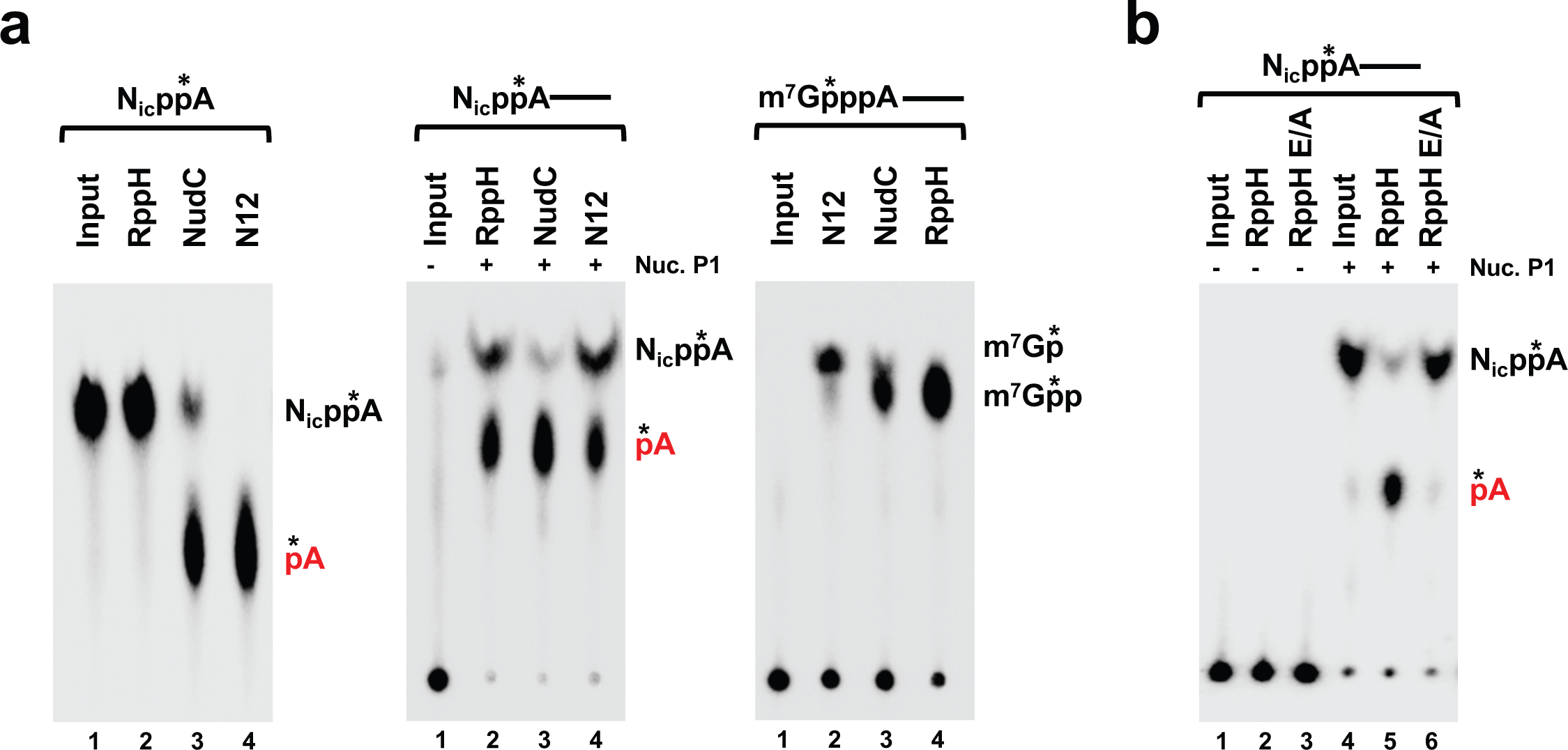
RppH has RNA deNADding activity *in vitro*. **(a)** *E. coli* RppH is a robust deNADding enzyme. *In vitro* decapping assays were carried out with 50 nM RppH, NudC or mouse Nudt12 and ^32^P-labeled substrates: free NAD (left panel), NAD-capped RNA (middle panel) and m^7^G-capped RNA (right panel). Labeling is as described in the legend to Fig. 1 and TLC was developed in 0.45 M (NH_4_)_2_SO_4_. **(b)** Catalytically inactive mutant RppHE53A (RppH E/A) lacks deNADding activity. *In vitro* deNADding assays were carried out with 100 nM *E. coli* RppH or RppH E/A proteins and ^32^P-labeled NAD-capped RNA substrate. TLC as developed in 0.5 M LiCl.

**Table 1.** deNADding Enzyme Classes

| <u>CLASS</u> | <u>ENZYME</u> | <u>SUBSTRATE</u> | <u>PRODUCT</u> |
| --- | --- | --- | --- |
| 1 | DXO<br>Dxo1<br>Rai1 | N<sub>ic</sub>ppA-RNA | N<sub>ic</sub>ppA<sub>OH</sub> + pRNA |
| 2 | NudC<br>Nudt12 | N<sub>ic</sub>ppA<br>or<br>N<sub>ic</sub>ppA-RNA | N<sub>ic</sub>p + pA<br>or<br>N<sub>ic</sub>p + pA-RNA |
| 3 | RppH | N<sub>ic</sub>ppA-RNA | N<sub>ic</sub>p + pA-RNA |

### Edc3 does not contain detectable deNADding activity *in vitro*

A stimulator of the Dcp2 mRNA decapping enzyme, enhancer of mRNA decapping 3 (Edc3) protein, is an NAD(H)-binding protein with potential hydrolytic activity on free NAD(H)^22^, which prompted us to test whether Edc3 has deNADding activity. As shown in Supplementary Fig. 1, under parameters where robust Nudt12 deNADding activity can be detected, human Edc3 protein failed to exhibit deNADding activity. Additionally, Dcp2, which lacks deNADding activity^7^, was also not stimulated by Edc3 to hydrolyze NAD-capped RNA (Supplementary Fig. 1). Our findings demonstrate mouse Nudt12 possesses deNADding activity *in vitro*, while the Edc3 NAD(H)-binding protein lacks deNADding activity.

### Structural insights into Nudt12 deNADding

To understand the molecular mechanism of Nudt12 deNADding, we determined the crystal structure of the catalytic domain of mouse Nudt12 (residues 126-462) in complex with AMP and 3 Mg^2+^ ions at 1.6 Å resolution (Supplementary Table 1). Nudt12 contains an Ankyrin repeat domain in the N-terminal region, which is absent in NudC (Fig. 3a). This domain is required for RNA deNADding activity, but is dispensable for NAD hydrolysis (Supplementary Fig. 2a), suggesting that the domain may contribute to binding the RNA body in the substrate.

**Figure 3.**
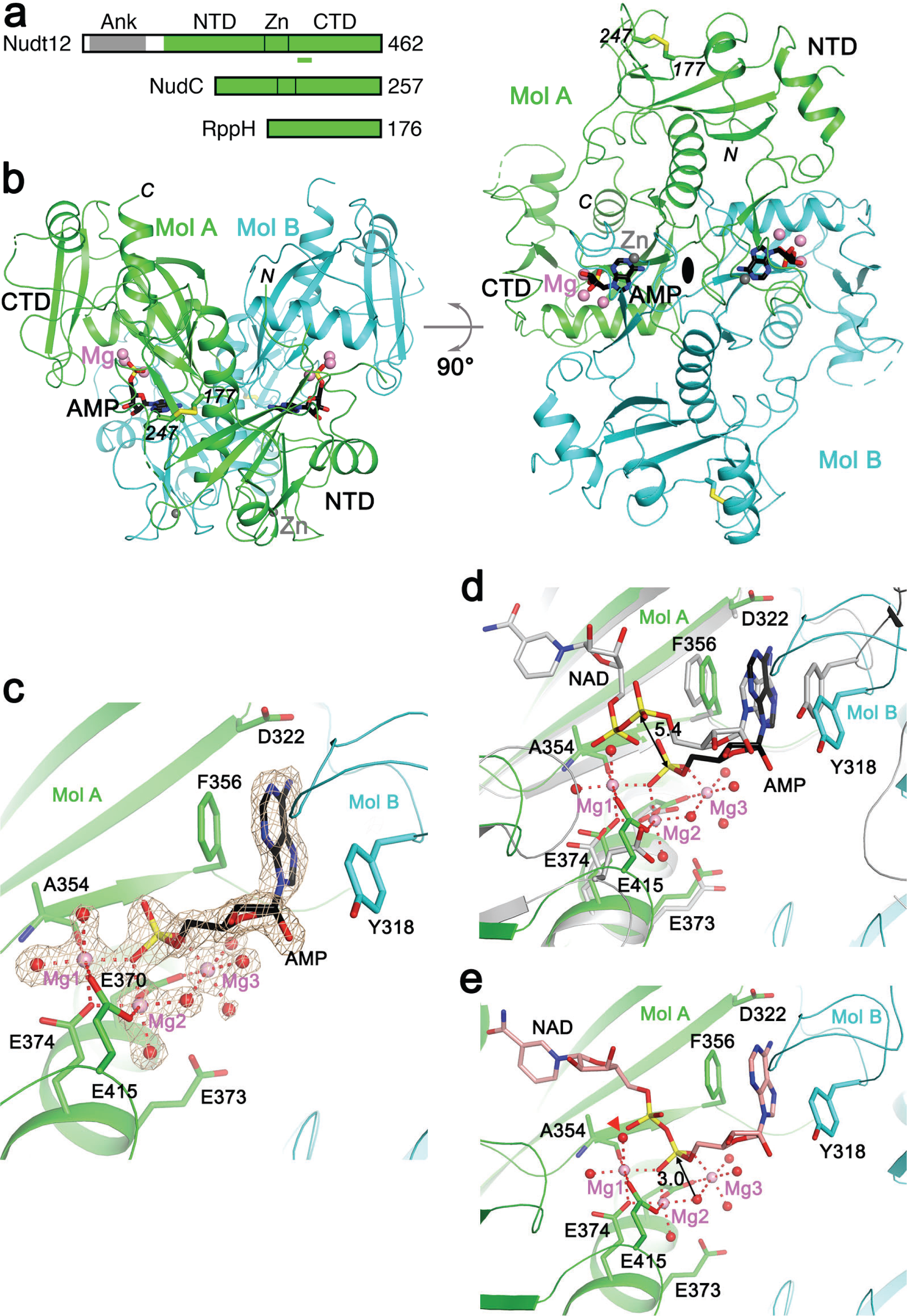
Crystal structure of mouse Nudt12 in complex with AMP and 3 Mg^2+^ ions. **(a)** Domain organization of mouse Nudt12 and *E. coli* NudC and RppH. The Ankyrin repeat domain of Nudt12 is indicated in gray, and catalytic domain in green. The N-terminal (NTD), Zn-binding (Zn) and C-terminal (CTD) sub-domains are also indicated. The conserved Nudix motif in Nudt12 is indicated with a bar (green). **(b)** Two views of the crystal structure at 1.6 Å resolution of mouse Nudt12 catalytic domain in complex with AMP and 3 Mg^2+^ ions. The two monomers are colored green and cyan, respectively. AMP is shown as stick models in black, Mg^2+^ ions as pink spheres, and Zn as gray spheres. A disulfide bond between residues 177 and 247, formed during crystallization, is indicated as stick models. **(c)** Detailed binding mode of AMP and Mg^2+^ ions in Nudt12. The coordination sphere of each metal ion is indicated with the red dashed lines. Water molecules are shown as red spheres. The omit F_o_–F_c_ electron density at 1.6 Å resolution for AMP, Mg^2+^ ions and their water ligands is shown in wheat color, contoured at 5 σ. **(d)** Comparison to the binding mode of NAD in NudC. Overlay of the structure of Nudt12 (in color) in complex with AMP (black) and three Mg^2+^ ions (pink) with that of NudC in complex with NAD (gray). The position of the phosphate of AMP is separated by 5.4 Å from the equivalent phosphate of NAD, indicated with the black arrow. **(e)** Molecular mechanism of the deNADding reaction. A model of the binding mode of NAD is shown (salmon color). The AMP portion of the model is identical to the crystal structure, and the NMN portion is based on that in the NudC complex. The nucleophilic attack by the bridging ligand of Mg2 and Mg3 on the phosphate is indicated by the black arrow, which initiates the deNADding reaction. The water molecule that is displaced in the NAD complex is indicated with the arrowhead (red).

The Nudt12 catalytic domain consists of an N-terminal sub-domain (NTD, residues 126-282), a zinc-binding motif (residues 283-318, where Zn is bound by four Cys side chains), and a C-terminal sub-domain (CTD, residues 319-462) (Fig. 3a). The conserved Nudix sequence elements are located in the CTD (Fig. 3a), although the NTD also has the Nudix fold, but with a more divergent structure. The catalytic domain forms a dimer with an extensive interface, involving contributions from all three sub-domains. The two monomers have essentially the same conformation, with rms distance of 0.46 Å for 303 aligned Cα atoms (Supplementary Fig. 2b). The catalytic domain shares 29% amino acid sequence identity with *E. coli* NudC. While the overall structure of Nudt12 is similar to that of NudC^23^, there are also substantial differences between them, especially for the NTD (Supplementary Figure 2b,c). The rms distance in Cα atoms is 2.0 Å for 226 aligned residues between the two proteins.

While we included NAD during crystallization, clear electron density was observed for AMP based on the crystallographic analysis (Fig. 3c), confirming that the catalytic domain of Nudt12 can hydrolyze NAD (Supplementary Fig. 2a). AMP is primarily bound to the CTD of one protomer (Fig. 3b). The adenine base is in the *syn* configuration, π-stacked with the side chain of Phe356 on one face and on the other face with that of Tyr318 in the loop connecting the zinc-binding motif and the CTD of the other protomer (Fig. 3c). There are no direct hydrogen-bonding interactions between the adenine base and Nudt12, although there are water-mediated interactions. The ribose hydroxyls of AMP are exposed to the surface, leaving substantial space to accommodate the RNA body in the NAD-capped RNA.

The phosphate group of AMP is involved in a large network of interactions with Nudt12, mediated through three Mg^2+^ ions (named Mg1, Mg2 and Mg3) (Fig. 3c). One of the terminal oxygen atoms of the phosphate is a bridging ligand to Mg1 and Mg2, while a second terminal oxygen atom is coordinated to Mg3. Three Glu residues in the Nudix motif are ligands of the Mg^2+^ ions, including Glu370 (bidentate interactions with Mg2 and Mg3), Glu374 (coordinated to Mg1), and Glu415 (bidentate interactions with Mg1 and Mg2). The main-chain carbonyl of Ala354 is coordinated to Mg1. Each Mg^2+^ ion is coordinated octahedrally, and seven water molecules or hydroxide ions complete the coordination spheres, one of which is a bridging ligand between Mg2 and Mg3 (Fig. 3c).

The binding mode of AMP in the current structure has significant differences compared to that of the AMP portion of NAD in the complex with NudC^11,23^ (Fig. 3d). Probably due to the absence of metal ions in the NudC complex, the phosphate group is pushed away from the Glu side chains in the active site, and the distance between the phosphorus atoms in the two structures is 5.4 Å (Fig. 3d). In addition, the adenine base is in the *anti* configuration in one NudC complex^23^ and both *anti* and *syn* configurations in the other NudC complex^11^. Many residues in the active site region have different conformations as well, especially the residue equivalent to the Glu415 ligand in Nudt12 (Fig. 3d). On the other hand, the NMN portion of NAD in NudC is located near the AMP in the current structure (Fig. 3d), and a model for the binding mode of NAD to Nudt12 could be built based on this (Fig. 3e). In this model, one of the terminal oxygen atoms on the phosphate in NMN can be directly coordinated to Mg1, displacing one of the water molecules in the current structure.

The model of the NAD complex provides clear molecular insights into the deNADing mechanism. The bridging ligand between Mg2 and Mg3 is located directly beneath the phosphate group of AMP, with a distance of 3.0 Å to the phosphorus atom (Fig. 3e). This ligand is likely a hydroxide ion, and initiates the deNADding reaction by attacking the phosphorus atom, causing breakage of the pyrophosphate bond. The terminal oxygen atom of AMP that is not coordinated to the Mg^2+^ ions is connected to the NMN of NAD and is the leaving group. There does not appear to be a general acid from the protein, in Nudt12 or NudC, that can stabilize the oxyanion of the leaving group, and this role might be served by a solvent molecule that becomes bound in the presence of NAD.

### Mammalian Nudt12 is a deNADding enzyme in cells

To test whether Nudt12 functions as a deNADding enzyme in cells, we asked whether the stability of NAD-capped RNA is altered in the absence of Nudt12. A collection of CRISPR/Cas9n directed individual or double knockout HEK293T cell lines comprised of a pool of three monoclonal knockout cell lines in the *Nudt12* gene (N12-KO), were generated and used (Fig. 4a). ^32^P uniformly labeled RNA transcripts containing either NAD or m^7^G at the 5′-end and 16 consecutive guanosine nucleosides (G_16_) at the 3′-end to minimize 3′-end decay^24^ were transfected into the respective knockout HEK293T cell lines. RNA was harvested at various times up to 4 h post transfection, resolved and detected. As shown in Fig. 4b, NAD-capped transcripts were more stable in N12-KO than in Con-KO cells (t_½_ = 4.1h *versus* 2.2h, p=3.45 × 10^−5^, ANOVA). In contrast, m^7^G-capped RNA showed similar half-lives in both cell line backgrounds (Fig. 4c), indicating that Nudt12 preferentially modulates the stability of NAD-capped RNAs transfected into HEK293T cells.

**Figure 4.**
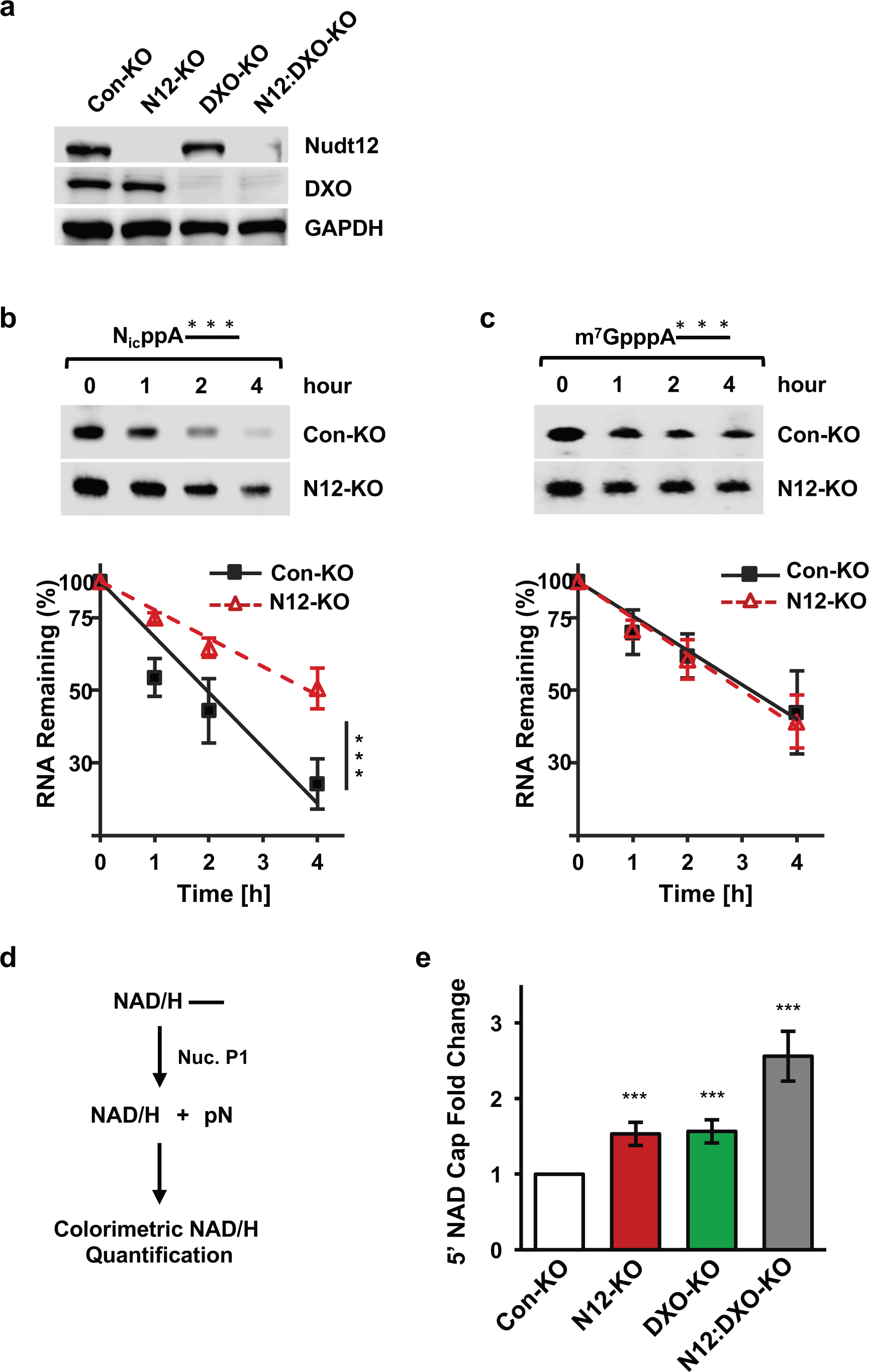
Mammalian Nudt12 is a deNADding enzyme in cells. **(a)** (Con-KO), Nudt12 knockout (N12-KO), DXO knockout (DXO-KO) or double knockout (N12:DXO-KO) cell lines. GAPDH was used as an internal control. **(b)** ^32^P uniformly labeled NAD-capped RNA, or **(c)** m^7^G-capped RNA were transfected into the indicated cell lines using Lipofectamine 3000. Untransfected RNAs were degraded with micrococcal nuclease (MN), and total RNA was isolated at the times denoted following micrococcal nuclease treatment. Remaining RNA was resolved on 8% denaturing PAGE, exposed to a PhosphorImager, quantitated and plotted. All data are derived from three independent experiments with ±SEM denoted by the error bars. **(d)** Schematic of NAD-cap detection and Quantitation (NAD-capQ) assay combining enzymatic properties of Nuclease P1 with a colorimetric NAD/NADH (NAD/H) Quantitation to detect total NAD and NADH levels. Background levels were derived by an identical assay that lacked Nuclease P1. **(e)** Total RNA from the indicated mammalian cells were subjected to NAD-capQ to detect levels of NAD-caps. Error bars represent ± SD. Statistical significance level was calculated by one-way ANOVA with a Dunnett’s multiple comparison posttest. p values are denoted by asterisks; (*) p < 0.05; (**) p < 0.01; (***) p < 0.001.

To directly determine whether Nudt12 contributes to expression of endogenous NAD-capped RNA in cells, we utilized an independent approach that detects NAD caps *en masse* termed NAD cap detection and Quantitation (NAD-capQ^18^). NAD-capQ combines the enzymatic properties of nuclease P1 to release intact NAD from the 5’ end of NAD-capped cellular RNA, with a colorimetric NAD quantitation to detect the released NAD (Fig. 4d). To determine whether levels of NAD-capped RNAs would increase in cells devoid of Nudt12, RNA from Con-KO or N12-KO cells were subjected to NAD-capQ. Consistent with Nudt12 functioning as a deNADding enzyme in cells, a 1.5-fold increase in total NAD-capped RNA was detected in N12-KO compared to the control. RNA from a pool of monoclonal cells disrupted for the *DXO* gene DXO-KO; ^7^; revealed a similar 1.5-fold increase in cellular NAD caps (Fig. 4e). Strikingly, a pool of monoclonal lines harboring a double knockout of both the *Nudt12* and *DXO* genes (N12:DXO-KO; Fig. 4a) resulted in an approximate doubling of NAD caps relative to the individual knockout backgrounds with 2.7 fold higher levels compared to Con-KO cells (Fig. 4e). Collectively, these data demonstrate that in addition to DXO^7^, Nudt12 is also a deNADding enzyme in cells and most significantly, they each appear to function on distinct NAD capped RNAs.

### Nudt12 preferentially targets a subset of mRNAs for deNADding

To directly determine whether Nudt12 contributes to the expression of specific NAD-capped RNAs in cells we utilized the NAD captureSeq approach which was successfully used to isolate NAD-capped RNAs in bacteria^2^, yeast^6^ and human^7^ cells. Briefly, NAD-capped RNAs are identified by the reaction of the 5’ end NAD with ADP ribosylcyclase (ADPRC) to generate an alkyne moiety amendable to click chemistry selective covalent incorporation of biotin and purification of the modified RNA with streptavidin^25^. NAD-captureSeq was carried out with RNAs from HEK293T WT or N12-KO cells. Selecting reference transcripts with at least a 2-fold increase in N12-KO compared with WT, at least 1 FPKM expression level in N12-KO and ≤5% false discovery rate (FDR) revealed 188 NAD-capped RNAs that were specifically enriched in the N12-KO cells (Supplementary Table 2) suggesting these are targets of Nudt12 deNADing. Importantly, subjecting the RNAs increased in Nudt12 knockout to gene ontology using high stringency (FDR ≤5%, ≥10 transcripts per term) identified five major pathways significantly enriched, three biological process (BP) terms and two cellular compartment (CC) terms (Fig. 5a). One of these relates to rRNA processing while the remaining are linked with mitochondrial metabolism or translation. Interestingly, while transcripts relating to rRNA processing are enriched, none of the 231 nuclear-encoded ribosomal protein gene transcripts was significantly different in N12-KO. Among the 150 mitochondrial-encoded ribosomal transcripts, only 10 are significantly enriched in N12-KO (listed in Supplementary Table 2, beginning with MRPL and MRPS). The mRNAs enriched in N12-KO are associated primarily with respiratory function, suggesting an interaction between cell metabolism and Nud12 deNADding of NAD-capped transcripts.

**Figure 5.**
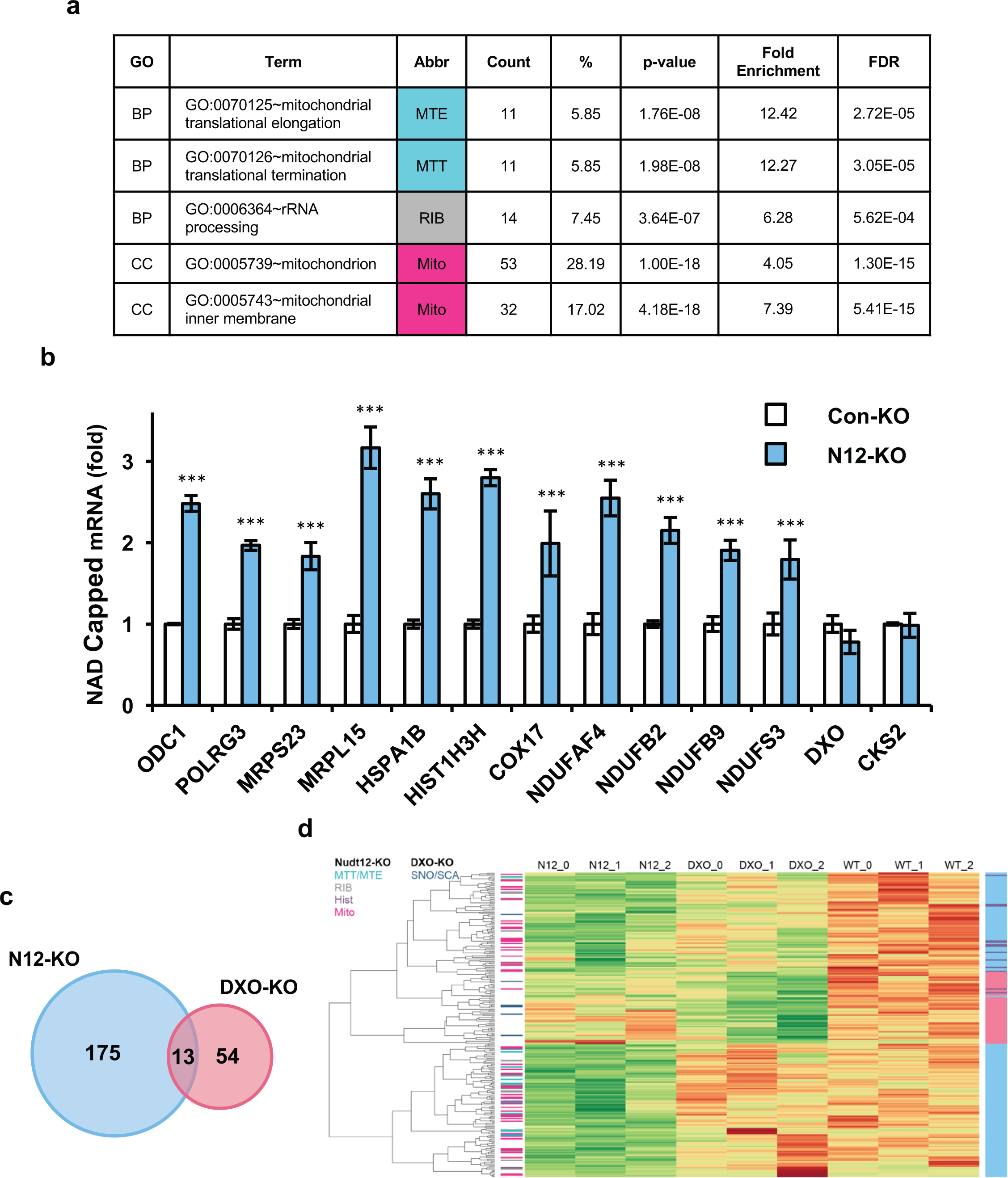
Nudt12 preferentially targets a subset of mRNAs for deNADding. **(a)** Top gene ontology (GO)-biological process (BP) and cellular component (CC) terms enriched with 188 genes increased in Nudt12-KO over control (FDR < 5%; > 2-fold increased, and > 1 FPKM in Nudt12-KO). GO terms were filtered for those with <5% FDR and at least 10 genes per term. **(b)** qRT-PCR validation of NAD-capped mRNAs in N12-KO cells. Randomly selected NAD-capped RNAs from the NAD-captureSeq were eluted from the beads, reverse transcribed, and detected with gene-specific primers. Data are presented relative to the HEK293T Con-KO cells and set to 1. CKS2, which is not responsive to Nudt12 in the NAD-captureSeq, was included as a negative control. Error bars represent ± SD. p values are denoted by asterisks; (***) p < 0.001 (Student’s t test). **(c)** A Venn Diagram of NAD-capped RNAs enriched in N12-KO and DXO-KO cells (≥2-fold, ≤ 5% FDR, > 1 FPKM in HEK293T WT cells). **(d)** Heatmap of mRNAs enriched in either N12-KO or DXO-KO. The color bar at left indicates enriched gene groups, either from top GO biological processes (Fig. 5a) or presence of major gene families as indicated (Hist: histones, SNO/SCA: snoRNAs or scaRNAs). The color bar at right indicates membership in components of Venn diagram (Fig. 5b). For each mRNA in the heatmap, green indicates relative enrichment, red indicates relative depletion, with expression normalized for each mRNA across all samples to indicate relative differences. Individual replicate samples from each group are indicated by a trailing number (“_0”, “_1”, etc.).

A subset of the NAD-captureSeq identified RNAs consisting of 5 RNAs implicated in oxidative phosphorylation and 7 randomly selected mRNAs were chosen for further validation by direct quantitative reverse-transcription (qRT)-PCR (Fig. 5b). We found that 11 out of 12 NAD-capped mRNAs identified from the NAD-captureSeq to be enriched in N12-KO cells were validated by direct qRT-PCR, while CKS2 mRNA, which was not a target of Nudt12 based on NAD captureSeq, was not changed in N12-KO compared to WT cells. Seven of the confirmed genes (COX17, MRPL15, MRPS23, NDUFA4, NDUFB2, NDUFB9, and NDUFS3; highlighted in Supplementary Table 2) were found in the enriched GO terms in Fig. 5a. These data further verify Nudt12 as a deNADding enzyme in cells and suggest a role for Nudt12 in nuclear-encoded NAD-capped transcripts encoding proteins with mitochondrial functions.

Applying similar data analysis to NAD-capped RNAs from DXO-KO cells^7^, we found that mRNAs enriched in the absence of Nudt12 were relatively distinct from those enriched when DXO was removed (Fig. 5c). Out of 188 unique RefSeq mRNAs enriched in N12-KO, only 13 overlap the 67 RefSeq mRNAs enriched by DXO-KO. Previous analyses identified many more DXO-KO-regulated transcripts from the same samples^7^, likely due to the inclusion of contrasting m^7^G-capped samples in the earlier analysis, which influenced the statistical evaluation. Interestingly, we found groups of NAD-capped RNA uniquely targeted by each enzyme (Fig. 5d). The hierarchical clustering of all 242 transcripts by replicate sample (Fig. 5d) indicates the distinct patterns of transcripts enriched in Nudt12 (light blue in right colorbar; matching Venn diagram) or DXO (coral in right colorbar). Highlighting the enriched GO terms in the left colorbar, the Nudt12-enriched GO terms are similarly distinct from a major DXO-KO category. Of the 175 genes enriched exclusively in N12-KO, two of the top GO-biological processes are linked to mitochondrial metabolism. In contrast, the most prominent class of NAD-capped RNAs elevated in DXO-KO cells, the small nucleolar RNAs (snoRNAs) and the related small Cajal body RNAs (scaRNAs)^7^ (Fig. 5d), were not elevated in NAD captureSeq from cells lacking Nudt12. NAD-capped mRNAs involved in oxidative phosphorylation were specifically enriched in N12-KO cells (Fig. 5). Collectively, these analyses revealed existence of Nudt12 and DXO responsive NAD-capped RNAs in mammalian cells and a modulation of distinct subset of mRNAs by each deNADding enzyme.

### Cellular exposure to environmental stress can impact NAD capping

Changes in growth conditions or exposure to environmental stress are often accompanied by changes in cellular NAD metabolism. In human cells, cellular NAD levels increase in response to energy stresses, such as glucose deprivation^26^, fasting^27,28^ and caloric restriction^29^, indicating that the nutrient state of a cell influences NAD concentrations. To assess whether physiological changes in cellular NAD can manifest altered NAD cap levels in cells, HEK293T cells were subjected to two distinct environmental stress conditions previously shown to affect cellular NAD metabolism: heat shock^30^ and glucose deprivation^26^. Cells were treated up to four hours with a 42°C heat shock or grown in glucose-free medium. Quantitation of NAD caps by NAD-capQ revealed ~2-fold increase in cells exposed to either heat shock or glucose starvation (Fig. 6a). The increase is not restricted to transformed cells. A similar glucose 2-fold increase in NAD cap levels was detected in glucose starved primary human foreskin fibroblast (HFF) cells (Fig. 6b). We conclude NAD-cap levels are dynamic and can increase in cells subjected to heat shock or glucose deprivation. These data suggest NAD capping provides a mechanism by which human cells can sense and respond to alterations in NAD metabolism.

**Figure 6.**
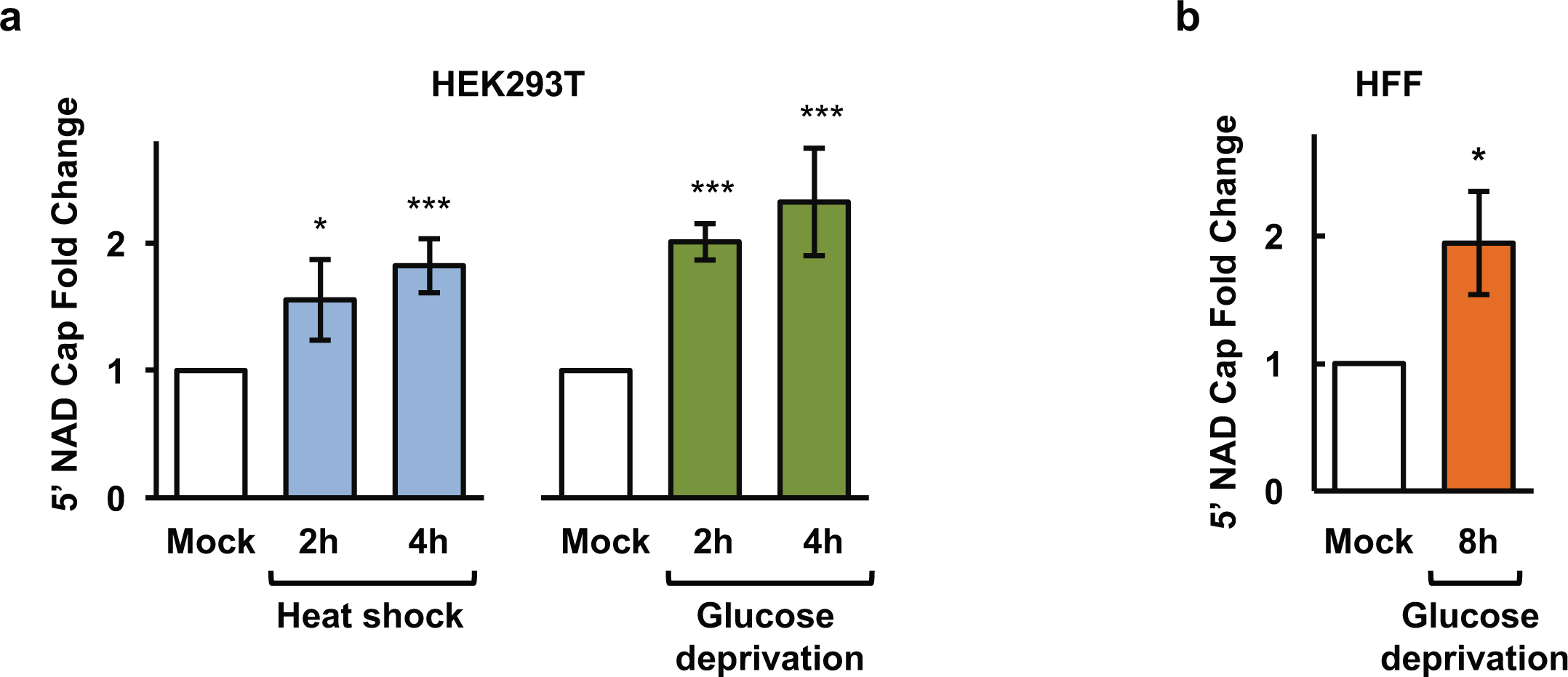
Cellular exposure to stress impacts NAD capping. **(a)** NAD-capped RNA levels were measured by NAD-capQ in HEK293T cells exposed to 42°C heat shock (left) or glucose deprivation (right) for the indicated times. **(b)** Same as **(a)** except from HFF cells grown in glucose depleted medium for indicated time. Graphs plot the fold-change in NAD-capped RNA values relative to the value observed in cells that were not subjected to stress. (Error bars represent ±SD, n ≥ 3; *, p < 0.05; ***, p < 0.001)

## DISCUSSION

Regulation of mRNA stability is an important post-transcriptional process that impacts eukaryotic gene expression. The presence of NAD caps in organisms ranging from *E. coli* to humans demonstrates the evolutionary conservation of the noncanonical NAD cap at the 5′ end of RNA transcripts and implies a function in RNA metabolism^16^. In this report, we demonstrate that, similar to the presence of multiple m^7^G cap decapping enzymes in cells^17^, mammalian cells also harbor more than just DXO as a deNADding enzyme. The Nudt12 Nudix hydrolase also functions as a deNADding enzyme in human cells. Nudt12 contains deNADding activity *in vitro* (Fig. 1), in human cells (Fig. 4b) and targets a specific group of mRNA for deNADding (Fig. 5). The crystal structure of mouse Nudt12 catalytic domain (residues 126-462) defines the binding mode of the AMP product and three coordinated magnesium ions. The structure also provides detailed insights into the molecular mechanism of the deNADding reaction (Fig. 3) and its distinct hydrolysis at the pyrophosphate linkage in Nudt12 to release NMN from NAD-capped RNA compared to the phosphodiester linkage in DXO that releases an intact NAD ^7^.

An important outcome of these studies was that Nudt12 is a deNADding enzyme that can promote the decay of NAD-capped RNAs in cells (Fig. 4) and it targets a subset of RNAs that are distinct from that of DXO (Fig. 5). The selective targeting of NAD-capped RNAs by the deNADding enzymes is supported by two complementary approaches. First, cells lacking either Nudt12 or DXO protein each contained an ~1.5-fold increase in total NAD-capped cellular RNA. Importantly, the difference was almost 3-fold in the double N12:DXO-KO cells (Fig. 4e) indicating that Nudt12 and DXO target different classes of NAD-capped RNAs for deNADding. Specificity of deNADding enzymes was further supported by identification of responsive target mRNAs. Specific NAD-capped RNAs, including histones and mitochondrial mRNAs are relatively enriched in N12-KO cells (Fig. 5d, green color) while depleted in DXO-KO cells (Fig. 5d, red color). Conversely, NAD-capped sno/scaRNAs are highly enriched in DXO-KO cells but not detected in cells lacking Nudt12.

The lack of a single mammalian deNADding enzyme that targets all NAD-capped RNAs in cells indicates deNADding enzymes preferentially target distinct mRNA subsets within different cellular pathways that similar to m^7^G cap decapping enzymes^17^. Intriguingly, 14 (19%) of the mRNAs encoding proteins within the mitochondrial electron transport chain contained NAD-capped versions that were elevated in Nudt12-KO cells. This is a much higher proportion than would be expected by chance alone (p=1.79 × 10-^16^; Hypergeometric distribution). Mitochondria are the main energy-generating organelles in cells, and these findings support a correlation between Nudt12 deNADding activity and cellular energetics.

An unanswered question remains as to how deNADding enzymes differentiate between different RNA substrates and preferentially deNAD specific RNAs. Two possibilities include selective direct binding to the 5’ end of specific RNAs or indirectly by recruitment through RNA-binding proteins. There is precedence for both approaches with the well characterized Dcp2 decapping enzyme. Dcp2 can bind a stem loop structure localized within 10 bases of the 5’ cap and preferentially decap the RNA^31,32^. Dcp2 can also selectively decap RNAs through recruitment by AU-rich RNA instability element binding proteins or proteins involved in nonsense mediated decay^33,34^. Whether Nudt12 and DXO utilize a direct or indirect mechanism to preferentially detect and deNAD specific NAD-capped RNAs remains to be determined.

An unexpected outcome of our studies was that the bacterial RppH Nudix protein also possesses deNADding activity, at least *in vitro* (Fig. 2a). The discrepancy with previous reports is not clear but may be a function of different *in vitro* parameters and/or different substrate RNA sequences employed in the two studies. RppH cleaves the NAD cap by hydrolyzing within the diphosphate bond similar to the activity observed with Nudt12 and the bacterial NudC Nudix proteins. These findings demonstrate that similar to mammalian cells, bacteria may also contain multiple deNADding enzymes. Importantly, although all three enzymes possess robust deNADding activity on NAD-capped transcripts *in vitro*, only Nudt12 and NudC, but not RppH, were capable of hydrolyzing free NAD (Fig. 2a), thus defining at least three distinct classes of deNADding enzymes (Table 1). Class I deNADding enzymes consist of the DXO family of proteins which remove an intact NAD from the 5’ end of NAD-capped RNAs^7^. Class II consists of proteins that can hydrolyze the pyrophosphate linkage of both free NAD as well as NAD at the 5’ end of an RNA and is represented by Nudt12 and NudC. Class III deNADding enzymes only function on NAD-capped RNA but not free NAD with RppH as the founding member.

Analysis of published structures of RppH provides some insight into its deNADding activity. The binding mode of the Mg^2+^ ions and the reaction mechanism of Nudt12 are generally similar to those observed for the *E. coli* RppH^35,36^ (Supplementary Figure 3a). One difference is that an Arg residue in RppH, Arg8, can stabilize the oxyanion of the leaving group. This Arg residue is equivalent to Asp322 in Nudt12 (Fig. 3c), and the observed position for its guanidinium group in RppH would clash with the side chain of Phe356 in Nudt12 (Supplementary Figure 3a), which is π-stacked with the adenine base (Fig. 3c). In addition, RppH is monomeric, covers only the CTD of Nudt12, and lacks the NTD and the zinc-binding motif (Fig. 3a).

The modeled binding mode for NAD in Nudt12 (Fig. 3e) is unlikely to be possible for RppH, as the nicotinamide portion would clash with the protein. However, a close comparison of the binding modes of pppRNA to RppH and NAD to Nudt12 suggests that a folded conformation of NAD (Supplementary Figure 3b), rather than the extended conformation as observed in Nudt12 and NudC (Fig. 3e), could fit in the RppH active site, where the adenine nucleotide of NAD would assume a conformation similar to that of the first nucleotide of pppRNA. Arg8 could also have cation-π interactions with the nicotinamide of NAD, in addition to its role in stabilizing the leaving group. The RNA body could assume the same binding mode for deNADding as for PPH activity (Supplementary Fig. 3b). Especially, the PPH activity of *E. coli* RppH has a preference for a guanine at the second position of the substrate^36^, suggesting that the deNADding activity would prefer a guanine nucleotide at the first position of the substrate RNA. Consistent with our biochemical data (Fig. 2), this model also suggests that RppH is only active toward NAD-RNA rather than NAD alone because the RNA body is important for substrate binding to RppH.

Another important finding from these studies was the demonstration that environmental stress and nutrient deprivation can both lead to elevation of NAD-capped RNA. NAD levels can fluctuate along with the metabolic state of a cell. Glucose restriction resulted in an increase of NAD-capped RNA in HEK293T and primary HFF cell lines (Fig. 6). These findings demonstrate that NAD capping can be modulated under stress conditions. Moreover, they potentially inform of a link between the redox state of a cell and RNA metabolism through a physiological link between NAD caps and environmental stress which will be an important area of future pursuit for the field.

## SUPPLEMENTAL INFORMATION

Methods and any associated references are available in the …….

## ACKNOWLEDGEMENTS

This work was supported by National Institutes of Health (NIH) grant GM118093 and S10OD012018 (L.T.), GM067005 and GM126488 (M.K.). We thank B.E. Nickels for helpful discussions and providing recombinant NudC. We thank K. Perry and R. Rajashankar for access to the NE-CAT 24-C beamline at the Advanced Photon Source. This work is based upon research conducted at the Northeastern Collaborative Access Team beamlines, funded by the NIH (P41 GM103403). The Pilatus 6M detector on 24-ID-C beam line is funded by a NIH-ORIP HEI grant (S10 RR029205). This research used resources of the Advanced Photon Source, a U.S. Department of Energy (DOE) Office of Science User Facility operated by Argonne National Laboratory under Contract No. DE-AC02-06CH11357. Computational resources were provided by the Office of Advanced Research Computing (OARC) at Rutgers, The State University of New Jersey, under the National Institutes of Health Grant No. S10OD012346.

## AUTHOR CONTRIBUTIONS

M.K., E.G.N., and L.T. designed the experiments. E.G.N. carried out all experiments unless otherwise indicated. X.J. and H.C. created N12 and N12:DXO CRISPR knockout cell lines. X.J. carried out the initial NAD captureSeq and the assays in Fig. 2. Y.W. and L.T. carried out the structural analysis and interpretations. R.P.H. carried out all bioinformatics analyses. E.G.N. M.K., L.T. and R.P.H. wrote the manuscript.

## COMPETING INTERESTS

The authors declare no competing interests.

## ONLINE METHODS

### Cell Culture and generation of Nudt12 and N12:DXO CRISPR knockout cell lines

Human embryonic kidney HEK293T cells were obtained from ATCC. Cells were cultured in DMEM medium (Thermo Fisher Scientific) supplemented with 10% fetal bovine serum (Atlanta Biologics), and antibiotics (100 units/ml penicillin and 100 µg/ml of streptomycin) under 5% CO_2_ at 37 °C. The three different monoclonal HEK293T cell lines harboring CRISPR-Cas9n double-nick generated distinct homozygous deletions within the *DXO* gene genomic region corresponding to its catalytic site have been reported previously^7^. Nudt12 knockout lines were similarly generated using CRISPR-Cas9n technology with two Nudt12-gRNAs (Supplementary Table 3) designed to target genomic regions of *Nudt12* corresponding to the catalytic site in exon 6. Nudt12:DXO double knockout lines were generated using the Nudt12 cell line and CRISPR-Cas9n technology with two DXO-gRNAs (Supplementary Table 3) designed to target genomic regions of *DXO* corresponding to the catalytic site in exon 4. The genomic modification was screened by PCR and confirmed by sequencing and a Western blot of the three mixed clones for N12-KO or N12:DXOKO is shown in Fig. 4a.

### Isolation and identification of NAD-capped RNAs

Total RNA from HEK293T, Con-KO, N12-KO, DXO-KO and N12:DXO-KO cell lines were isolated with TRIzol reagent (Thermo Fisher Scientific). NAD-capped RNAs were isolated using the NAD-RNA capture protocol^2^ with minor modification. NAD-capture was carried out with 50 µg total RNA treated with 10 µl 4-pentyn-1-ol (Sigma-Aldrich) and 3 U Adenosine diphosphate-ribosylcyclase (ADPRC) in 100 µl reaction containing 50 mM HEPES, 5 mM MgCl_2_ (pH 7) and 40 U RNasin^®^ Ribonuclease Inhibitor (Promega) at 37 °C 60 min. The reaction was stopped with phenol/chloroform extraction and RNAs were precipitated with ethanol in the presence of 2 M (final concentration) ammonium acetate. RNAs with a 5’ end NAD were biotinylated by treatment with a copper-catalyzed azide-alkyne cycloaddition (CuAAC) reaction by incubating with 250 µM biotin-PEG3-azide, freshly mixed 1 mM CuSO4, 0.5 mM THPTA, 2 mM Sodium Ascorbate in 100 µl reaction with 50 mM HEPES, 5 mM MgCl_2_ (pH 7) and 40 U RNasin^®^ Ribonuclease Inhibitor (Promega) at 30 °C for 30 min. RNA was precipitated with ethanol in the presence of 2 mM EDTA and 2 M (final concentration) ammonium acetate. Biotinylated NAD capped RNAs were captured by binding to 20 µl streptavidin magnet beads (Nvigen) at room temperature with gentle rocking for 1 hour in 100 µl binding buffer (1 M NaCl, 10 mM HEPES (pH 7.5) and 5 mM EDTA). The beads were washed three times with wash buffer (8 mM Urea, 50 mM Tris-HCl (pH 7.4), 0.1% IPEGAL) and two washes with nuclease free H_2_O. Biotinylated NAD-capped RNAs were eluted by resuspending beads in 100 µl H_2_O and heating to 95 °C for 5 min and RNA precipitated with ethanol in the presence of 2 M ammonium acetate. Captured NAD-RNAs were reverse transcribed into cDNA and synthesized into dsDNA with the SMARTer® Universal Low Input RNA Kit for Sequencing (Clontech Laboratories, Inc.) following the manufacturer’s protocol. NAD-capture for the qRT-PCR validation studies in Figure 5 were carried out similarly except, 50 ng of *in vitro* transcribed NAD-F.Luc-RNA was included as an internal control for normalization.

### *In vitro* Transcription of NAD-Capped RNAs

RNAs containing different cap structures were synthesized by *in vitro* transcription of plasmid ɸ2.5A-CA2-G16 containing the T7 ɸ2.5 promoter^37^ that retained the first adenosine at the transcription start site. All other adenosines within the RNA were replaced by uracils to maintain only one adenosine per RNA and a G-tract of 16 guanosines (G16) was present at the 3′end^7^. *In vitro* transcription was carried out at 37 °C for 90 min in 50 µl reaction containing 400 ng PCR-generated DNA template, 10 µl 5x transcription buffer, 5 µl DTT (10 mM), 5 µl BSA (10 µg/µl), 5 µl rNTP Mix (5 mM), 2 µl T7 RNA polymerase (Promega) and 1 µl RNasin Ribonuclease Inhibitor (40 U/µl, Promega). To generate NAD-capped RNA, rATP was replaced with ^32^P-NAD or NAD in the reaction to generate ^32^P-NAD-cap labeled or unlabeled NAD-capped RNAs, respectively. Transcription in the presence of [α-^32^P]GTP and 0.5 mM m^7^GpppA cap analog (New England BioLabs) minus rATP in the reaction was carried out to produce ^32^P-uniformly-labeled 5’-end N7-methylated RNA.

### RNA in vitro Decapping and deNADding Assays

^32^P-NAD-cap labeled or ^32^P-m^7^G-capped RNAs were incubated with the indicated amount of recombinant proteins in decapping buffer containing 10 mM Tris-HCl pH 7.5, 100 mM KCl, 2 mM DTT, 2 mM MgCl_2_ and 2 mM MnCl_2_ as previously described^38^ and incubated at 37 °C for 30 min. Reactions were stopped with 30 mM EDTA. For reactions involving Nuclease P1 treatment, reactions were first extracted with phenol followed by chloroform and 1U of Nuclease P1 (Sigma-Aldrich) was added. The reactions were incubate at 37 °C for 30 min. Decapping products were resolved by PEI-cellulose TLC plates (Sigma-Aldrich) and developed in 0.5 M LiCl or 0.45 M (NH_4_)_2_SO_4_ in a TLC chamber at room temperature^39^. Reaction products were visualized and quantitated with a Molecular Dynamics PhosphorImager (Storm860) with ImageQuant-5 software.

### RNA *in vivo* decay assay

Indicated cells (8 × 10^6^) were seeded in 100 mm plates one day prior to the experiment. 0.5 µg ^32^P-uniformly-labeled RNA was mixed with 10 µl P3000 reagent in 250 µl Opti-MEM medium and 15 µl Lipofectamine 3000 was diluted in 250 µl Opti-MEM medium (Thermo Fisher Scientific). The two solutions were mixed and incubated at room temperature for 15 min followed by addition of 3.5 ml Opti-MEM to the RNA-Lipofectamine 3000 complexes and applied to the cell culture plates for 30 min. The cells were washed with Opti-MEM and removed from the dish. Untransfected RNAs were degraded by Micrococcal Nuclease (New England Biolabs) in the presence of 5 mM CaCl_2_ for 30 min at 37 °C. The transfected cells were equally aliquoted into 60 mm cell culture plates and cultured at 37 °C. Cells were harvested at 0, 1, 2 and 4 hour time points after Micrococcal Nuclease treatment. Total RNA was isolated with TRIzol reagent and equal amounts of total RNA from the different time points were fractionated by 8% polyacrylamide gel with 7 M urea. The remaining ^32^P-labeled RNA was analyzed by PhosphorImager.

### RNA-Seq analysis

RNA was prepared, enriched by NAD-Capture, and sequenced as described^7^. Sequences were aligned with human hg19 genome (GRCh37) using the UCSC reference transcript map using Tophat ^40^ and the resulting binary sequence alignment/map format (BAM) files were sorted, duplicates were removed using Picard (Broad Institute; http://broadinstitute.github.io/picard), and transcripts were quantified using Cuffdiff ^40^. Results were exported to the CummeRbund package^40^ in R/Bioconductor. Only transcripts with expression levels of FPKM (fragments per kilobase of transcript length per million aligned reads) greater than 1.0 in the knockout condition were considered to be detectable. Sequencing data have been deposited in the Gene Expression Omnibus (GEO) database (accession no. GSE90884 (DXO-KO) and GSE110801 (N12-KO).

### Protein Expression, Purification and Crystallization

NudC recombinant protein was kindly provided by Bryce Nickels (Waksman Institute of Microbiology, Rutgers University). Nudt12, Nudt13 and RppH recombinant proteins carrying an N-terminal His-tag were expressed from plasmids pET28a-Nudt12, pET28a-Nudt13 and pET28a-RppH described previously^19^.

Residues 126-462 of wild-type mouse Nudt12 were sub-cloned into the pET28a vector (Novagen), and the recombinant protein contained an N-terminal His-tag. This purified wild-type protein did not produce any crystallization hits. The E219A/E220A/E221A triple mutation was introduced using Transfer-PCR method ^41^. The mutant protein was expressed in *Escherichia coli* BL21(DE3) Star cells at 20 °C for 18 h. The cells were lysed by sonication in a buffer containing 20 mM Tris (pH 8.0), 250 mM NaCl, and 5% (v/v) glycerol. The lysate was loaded onto an Ni-NTA (Qiagen) column. The eluted protein was treated overnight with thrombin at 4 °C to remove the His-tag and was further purified by gel filtration chromatography (Sephacryl S-300; GE healthcare). The purified protein was concentrated to 23 mg/ml in a solution containing 20 mM Tris (pH 8.0), 200 mM NaCl, and 5 mM DTT before being flash-frozen in liquid nitrogen and stored at –80 °C.

For crystallization, the above triple mutate protein at 8 mg/ml concentration was incubated with 1.5 mM NAD (Sigma) in a buffer containing 20 mM Tris (pH 8.0), 200 mM NaCl, 5 mM DTT, and 2 mM MgCl_2_ at 4 °C for 30 min. Crystals were obtained at 20 °C using the sitting-drop vapor-diffusion method. The reservoir solution contained 0.1 M HEPES (pH 7.5) and 24% (w/v) PEG2000MME. The crystals were cryo-protected by the reservoir solution supplemented with 15% (v/v) ethylene glycol and were flash-frozen in liquid nitrogen for data collection at 100K.

### Data Collection and Structure Determination

X-ray diffraction data were collected using the Pilatus-6M detector at the Advanced Photon Source (APS) beamline 24-ID-C. The X-ray wavelength was 0.9791 Å. The diffraction images were processed and scaled using the XDS program ^42^. The crystal belonged to space group *P*1 with unit cell dimensions of *a* = 56.2 Å, *b* = 58.6 Å, *c* = 61.7 Å. There is a Nudt12 homodimer in the crystallographic asymmetric unit.

The catalytic domain of Nudt12 shares 29% sequence identity with NudC, and a molecular replacement solution could readily be found using the structure of NudC as the search model^23^ with the program Phaser^43^. However, the resulting electron density map was not of sufficient quality to rebuild the model. To obtain separate phase information, the CTD of Nudt12 was located using the molecular replacement method. The phase information from this model was used to calculate an anomalous difference electron density map, which clearly revealed the positions of the two zinc atoms (with 11α peak heights), even though the diffraction data were collected far above the zinc absorption edge. Phase information from anomalous scattering was combined with that from the model, and the CTD could be rebuilt automatically using the resulting map with PHENIX^44^. Several β-strands in the NTD and segments near the zinc could be built manually with the program Coot^45^ and were included in the subsequent structure refinement with PHENIX, which led to an improved 2F_o_–F_c_map. The entire structure was obtained after several rounds of manual model building followed by refinement. The crystallographic information is summarized in Supplementary Table 1.

### RNA isolation, reverse transcription, and real-time qRT-PCR

Total cellular RNA was harvested with TRIzol Reagent (Thermo Fisher Scientific) and treated with DNase (Promega) according to the manufacturers’ protocols. Reverse transcription was performed on 2 µg of RNA in 20-µl reaction mixtures with M-MLV reverse transcriptase, random hexamers, and oligo(dT) (Promega) according to the manufacturer**’**s instructions. qRT-PCR was performed with the primers listed in a Supplementary Table 4. qRT-PCR was carried out on Rotor-Gene 3000 (Corbett Research) with iTaq SYBR Green Supermix (Bio-Rad Laboratories). Relative mRNA levels were normalized to 18S rRNA and calculated as described in User Bulletin No. 2 for the ABI Prism 7700 Sequence Detection System.

### NAD-cap detection and Quantitation

HEK293T cells (6 × 10^6^) were seeded in 100 mm plates a day before the experiment. Cells were ~ 80% confluent at the time of protein or RNA extraction. To measure NAD-cap amounts on RNA, total cellular RNA was extracted with TRIzol Reagent according to the manufacture’s protocol (Thermo Fisher Scientific). To maximize removal of potential residual free NAD copurifying with the RNA, isolated RNA was dissolved in 10 mM Tris-HCl (pH 7.5) containing 2 M urea. Samples were incubated 2 min at 65 °C and immediately precipitated with isopropanol in the presence of 2 M (final concentration) ammonium acetate. Fifty micrograms of total RNA was subjected to NAD-Cap Detection and Quantitation (NAD-capQ) assay as described^18^. Briefly, total RNA was digested with 1 U of Nuclease P1 (Sigma-Aldrich) in 20 µl of 10 mM Tris (pH 7.0), 20 µM ZnCl_2_ at 37 °C for 20 min to release 5’ end NAD. The control samples were prepared by incubating 50 µg of RNA treated with the same reaction condition including 10% glycerol used to dissolve the enzyme, but lacked Nuclease P1. After the reaction, 30 µl of NAD/H Extraction Buffer (NAD/H Quantitation Kit, Sigma-Aldrich) was added to each sample. In the second step, 50 µl samples were used to perform the colorimetric assay according to the manufacture’s protocol (NAD/H Quantitation Kit, Sigma-Aldrich).

### Western blot analysis

Cells were lysed in phosphate-buffered saline containing 0.5% IGEPAL CA-630 (Sigma-Aldrich), protease inhibitors (Roche Applied Science) and sonicated. Equal protein amounts of the different samples were separated by Bolt 4-12% Bis-Tris Plus Gel (Thermo Fisher Scientific) and transferred to nitrocellulose membranes (Bio-Rad). The blots were incubated with primary antibodies in phosphate-buffered saline buffer supplemented with 5% BSA (Sigma-Aldrich), then with secondary antibodies conjugated to horseradish peroxidase. Proteins were detected using the ECL western blotting substrate (Thermo Fisher Scientific).

## QUANTIFICATION AND STATISTICAL ANALYSIS

mRNA half-lives of reporter mRNAs were obtained by constructing linear models of the ln[remaining RNA] as a function of time, and then the half-life calculated as the ln[2]/slope^46^. All data are presented ± the 95% confidence interval and p values were calculated from the linear model using R^47,48^.

Following alignment with genome sequences, sequencing reads were assembled with the UCSC hg19 reference transcript map using Cufflinks, allowing a direct comparison of expression levels by Cuffdiff. Results were exported to the CummeRbund package^40^ in R/Bioconductor^47,48^. Only transcripts with expression levels of FPKM (fragments per kilobase of transcript length per million aligned reads) greater than 1.0 in at least one condition were considered to be expressed.

## DATA AVAILABILITY

The data that support the findings of this study are available from the corresponding author upon reasonable request. Sequencing data have been deposited in the Gene Expression Omnibus (GEO) database (accession no. GSE90884 (DXO-KO) and GSE110801 (N12-KO). The atomic coordinates have been deposited at the Protein Data Bank (accession number to be provided at proof stage).

---

**Figure S1.**
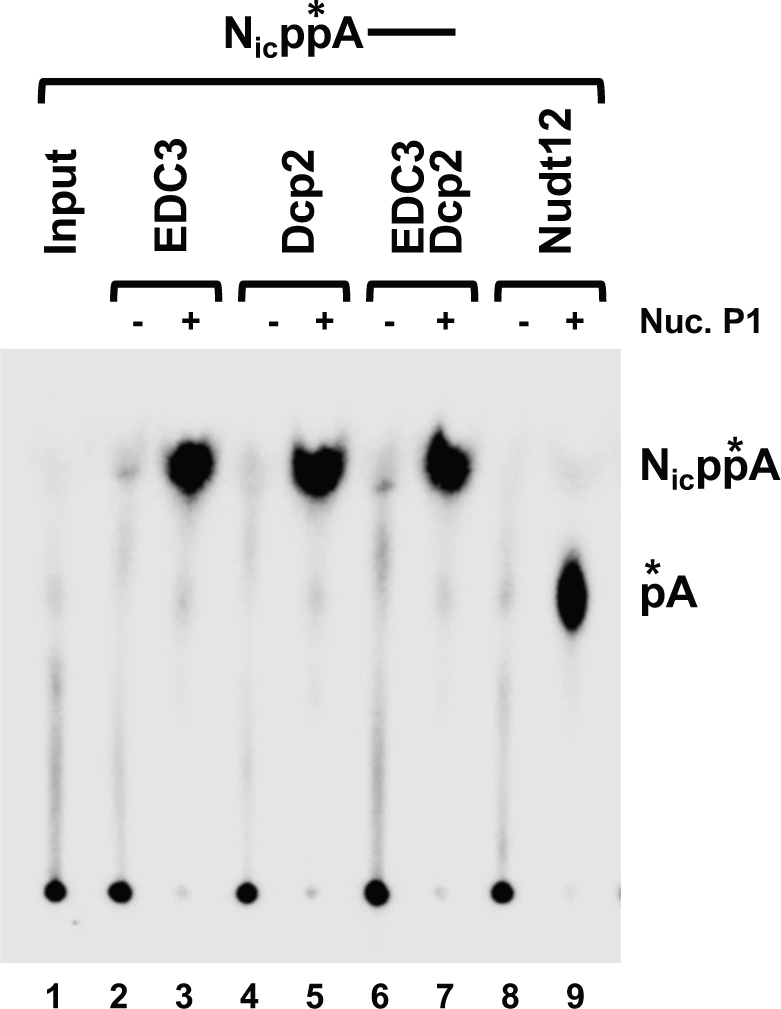
Edc3 NAD(H)-binding protein lacks deNADding activity. *In vitro* deNADding assays were carried out with 100 nM Edc3, Dcp2 or Nudt12 proteins and N_ic_pp*A-capped RNA as a substrate. DeNADding products were resolved by (PEI)-cellulose TLC plates developed in 0.45 M (NH_4_)_2_SO_4_.

**Figure S2.**
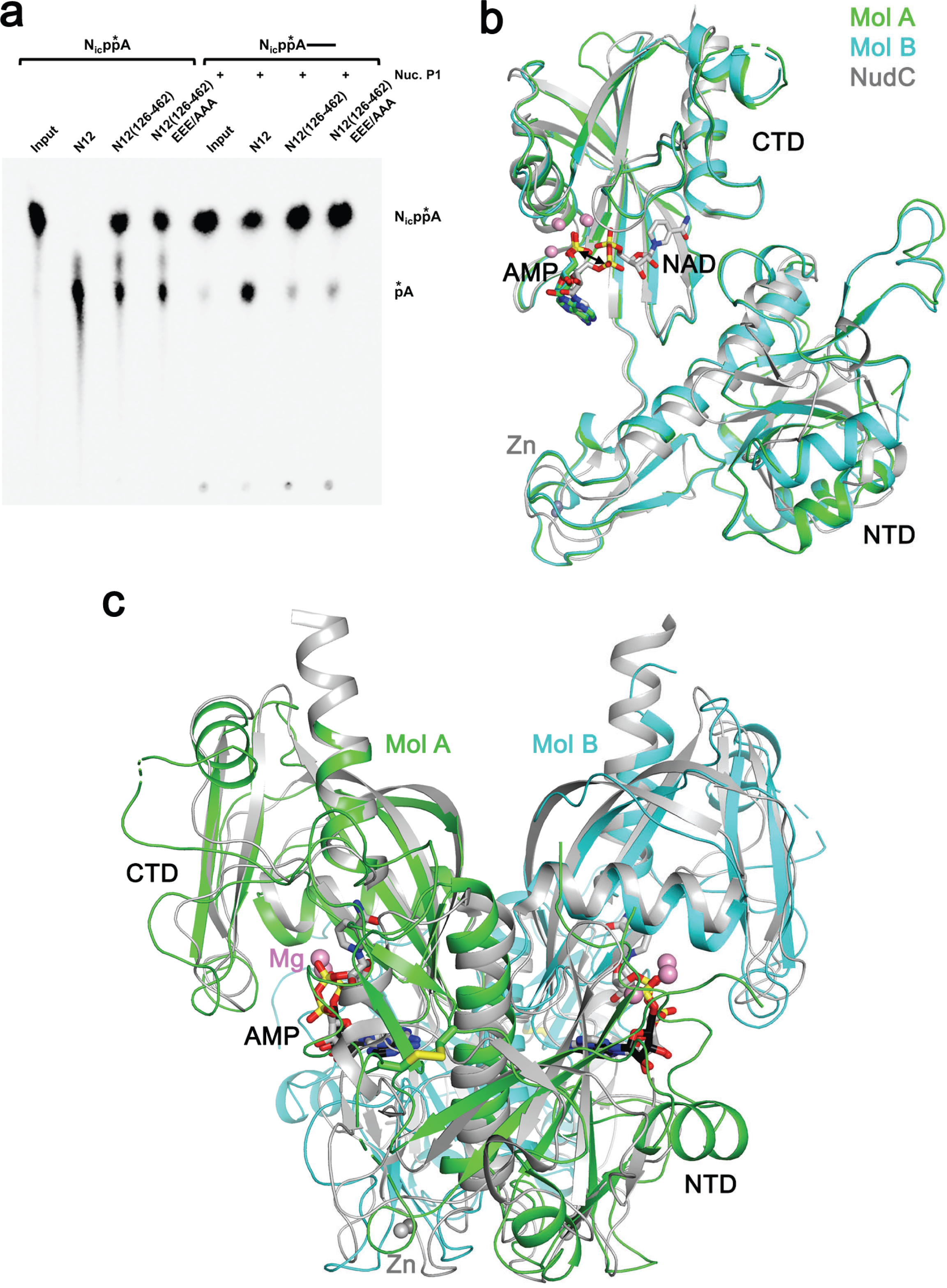
**(a)** The Ankyrin repeat domain of Nudt12 is required for deNADding activity on NAD-capped RNA but dispensable for NAD hydrolysis. The sample used for structural studies here, residues 126-462, can hydrolyze NAD. The EEE/AAA mutation, E219A/E220A/E221A, was introduced for aiding crystallization, and it does affect the catalytic activity. **(b)** Overlay of the structures of the Nudt12 monomers in complex with AMP (green and cyan) and Mg^2+^ (pink spheres) as well as NudC in complex with NAD (gray). The NTD shows substantial structural differences between Nudt12 and NudC. **(c)** Overlay of the structures of the Nudt12 dimer (green and cyan for the two monomers) and the NudC dimer (gray). Substantial structural differences are observed, especially for the NTD.

**Figure S3.**
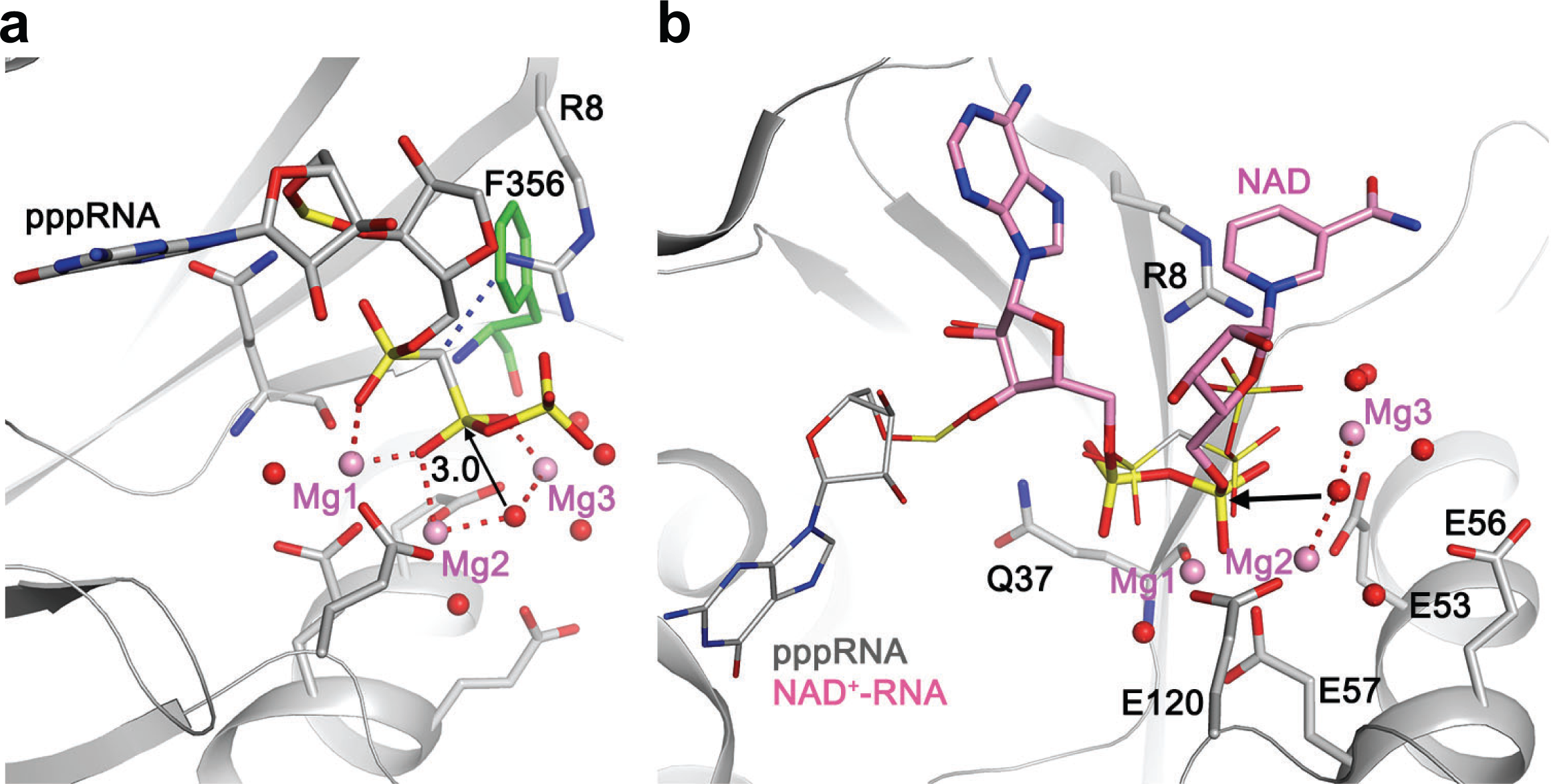
**(a)** Comparison to the substrate binding mode and reaction mechanism of RppH. The structure of RppH in complex with pppRNA is shown^36^, in the same orientation as that for Nudt12 (Fig. 2e). Residue Phe356 in Nudt12 is shown (green), clashing with the side chain of Arg8 in RppH. **(b)** A model for the binding mode of NAD to RppH and the molecular mechanism for its deNADding activity. The AMP portion of NAD (pink) is based on that in Nudt12, and the nicotinamide mononucleotide portion is based on the first nucleotide of pppRNA (gray) in RppH. The amide group of nicotinamide could be recognized by hydrogen-bonding interactions (dashed lines in red). Arg8 could have cation-π interactions with the adenine base as well as stabilize the leaving group.

**Supplementary Table 1.** Summary of crystallographic information. 1. Numbers in parentheses are for the highest resolution shell.

| **Data collection** |  |
| --- | --- |
| Space group | *P*1 |
| Cell dimensions |  |
| *a*, *b*, *c* (Å) | 56.2, 58.6, 61.7 |
| $\alpha$, $\beta$, $\gamma$ (°) | 102.4, 115.2, 104.7 |
| Resolution (Å)<sup>1</sup> | 53-1.6 (1.69-1.6) |
| *R*<sub>merge</sub> (%) | 4.3 (65.3) |
| CC<sub>1/2</sub> | 0.999 (0.716) |
| *I*/σ*I* | 11.5 (1.2) |
| Completeness | 89.6 (76.3) |
| No. of observations | 214,665 (27,577) |
| Redundancy | 2.8 (2.6) |
| **Refinement** |  |
| Resolution (Å) | 53-1.6 |
| No. of reflections | 76,425 |
| *R*<sub>work</sub> | 18.0 |
| *R*<sub>free</sub> | 21.9 |
| No. of atoms |  |
| Protein | 4,969 |
| Ligand/Ion | 54 |
| Water | 290 |
| B-factors |  |
| Protein | 39.3 |
| Ligand/Ion | 28.4 |
| Water | 38.8 |
| R.m.s. deviations |  |
| Bond lengths (Å) | 0.015 |
| Bond angles (°) | 1.4 |

**Supplementary Table 2.**
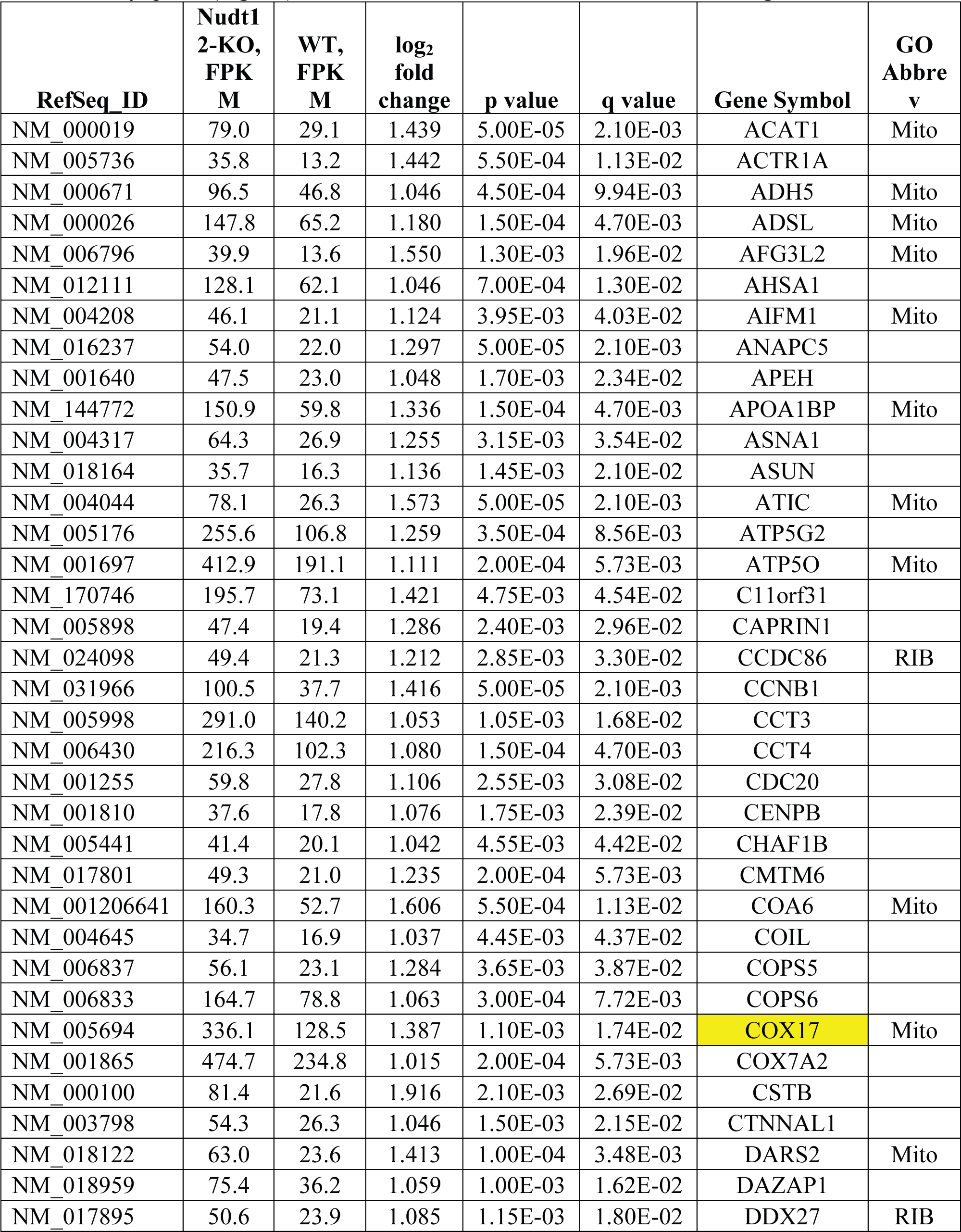
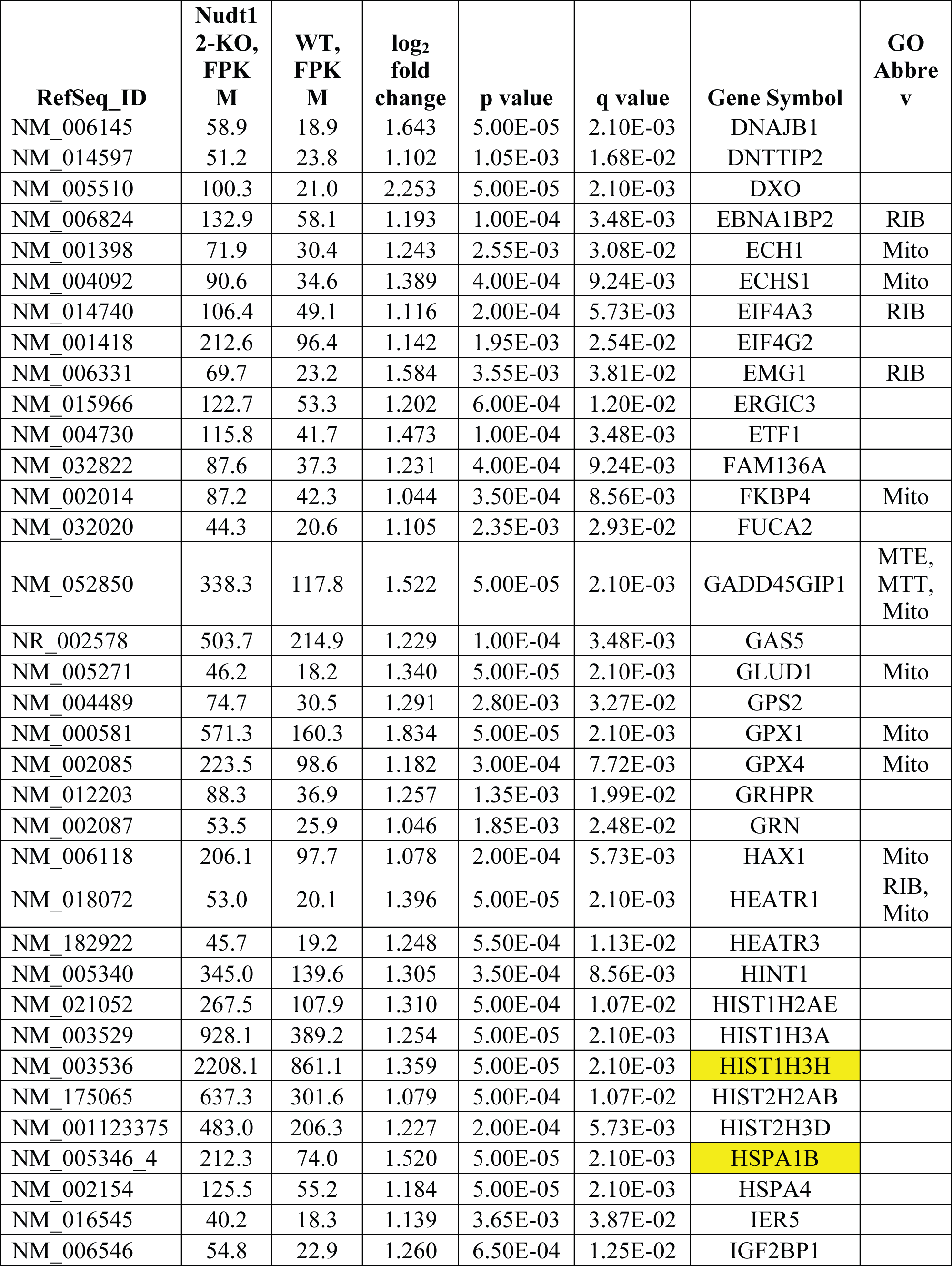
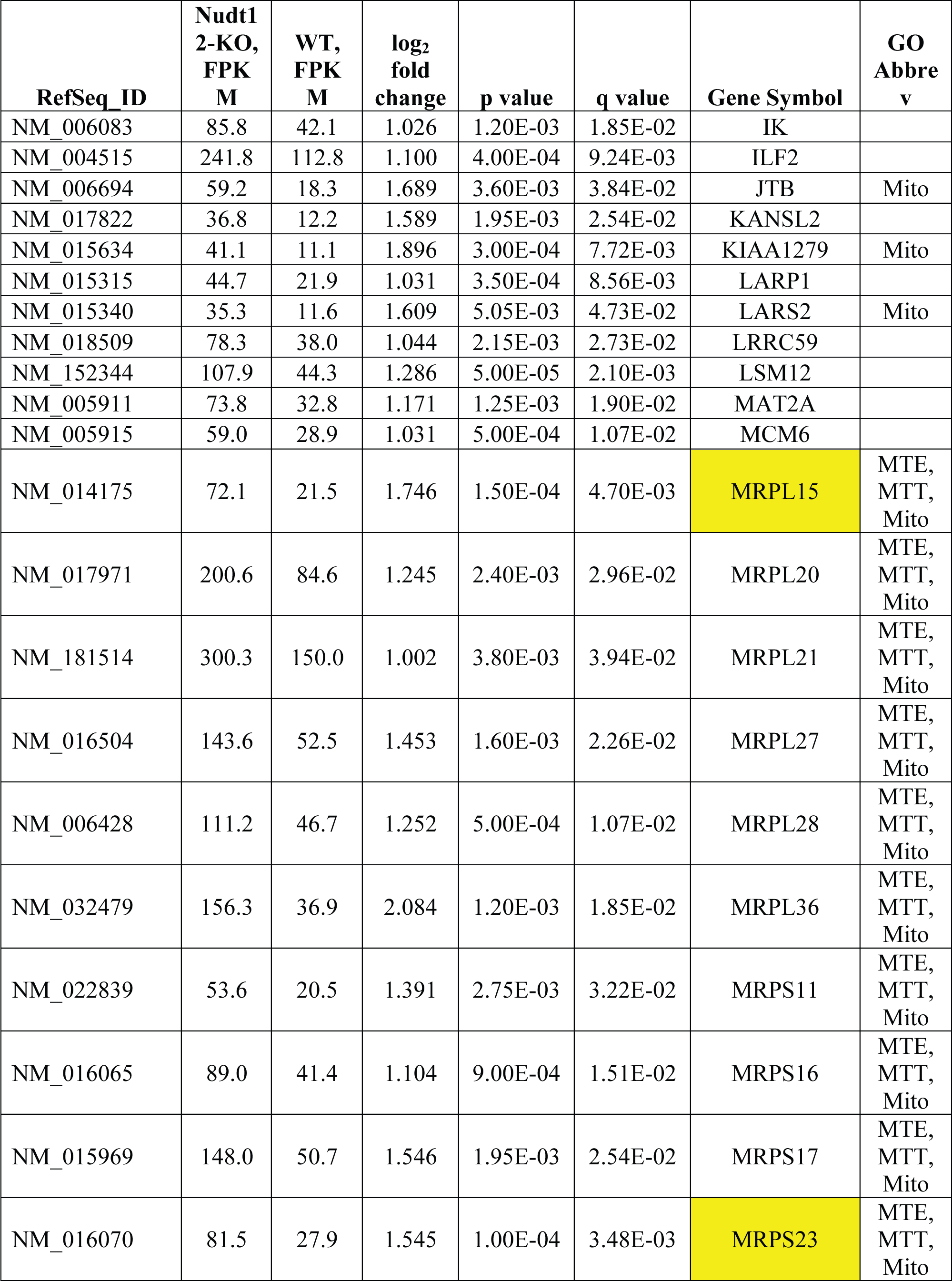
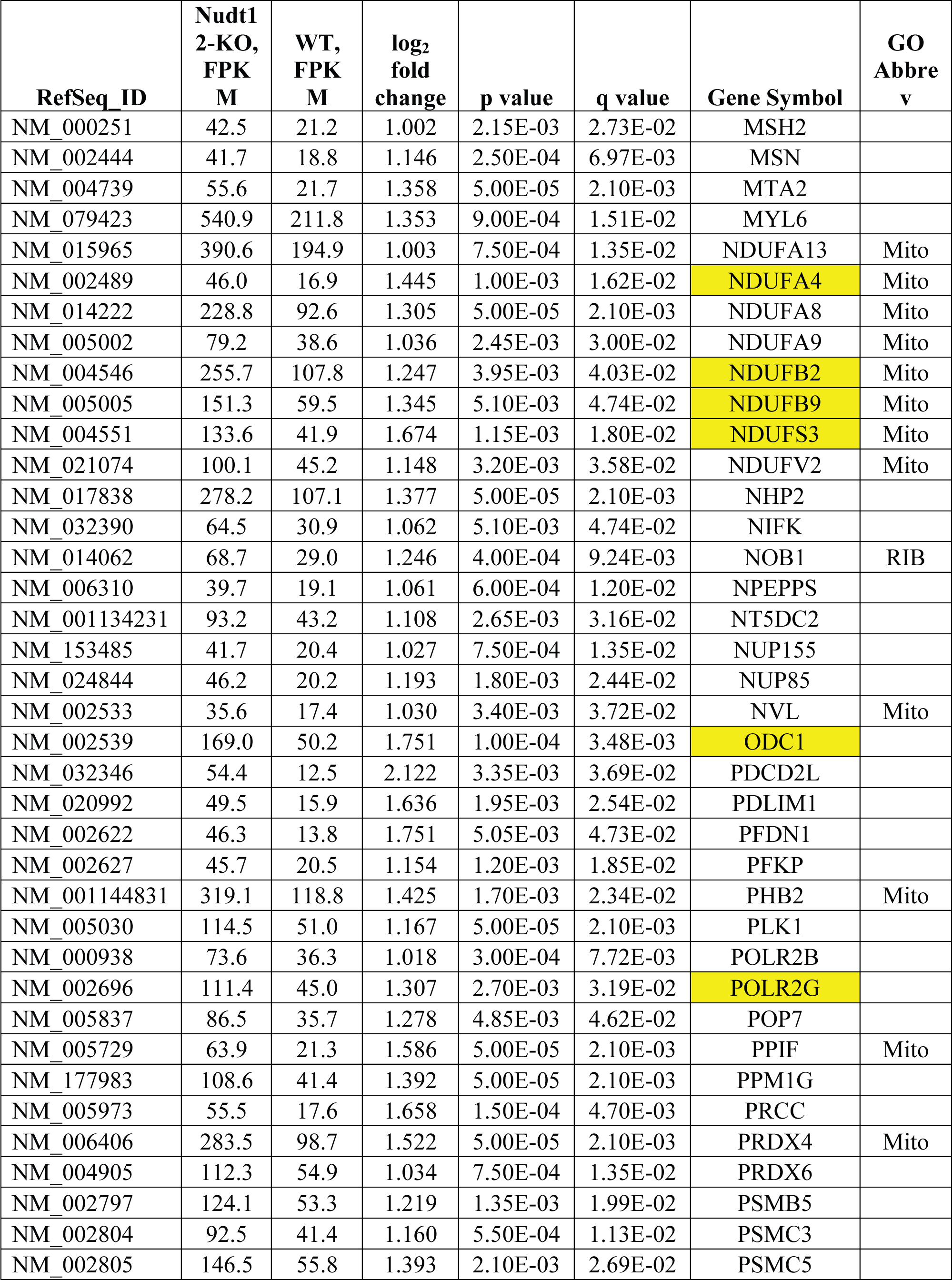
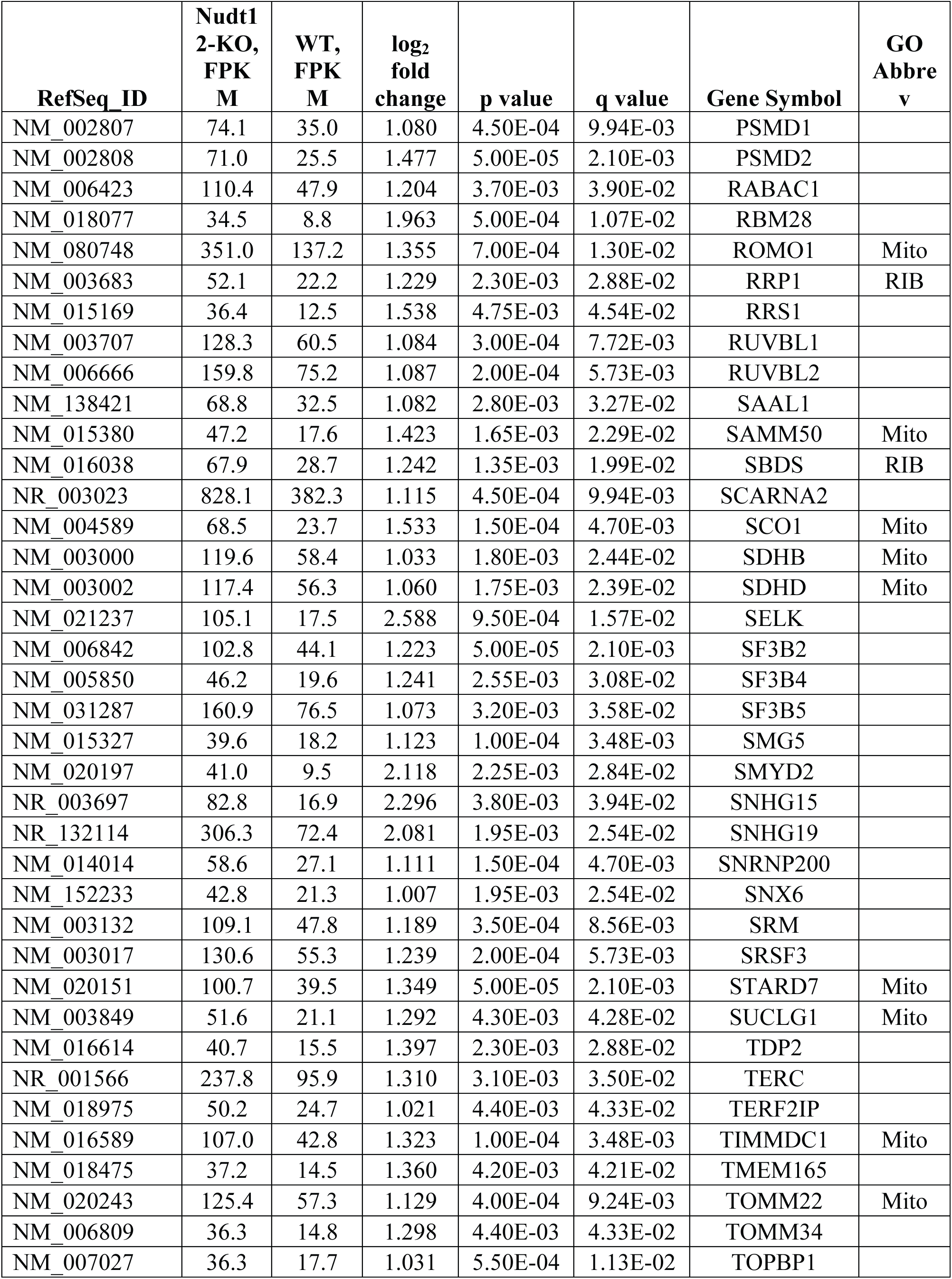
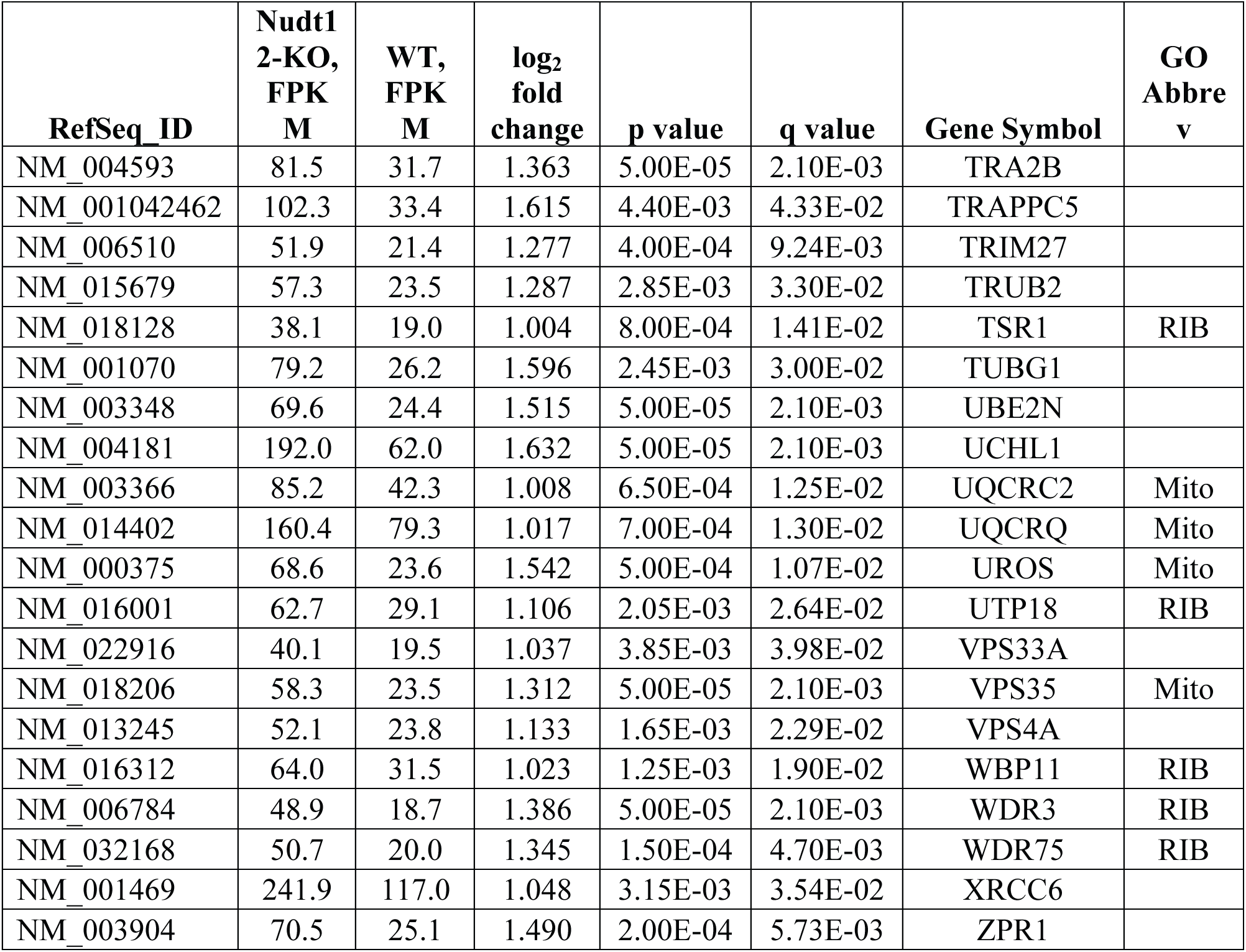
Genes included in enriched GO terms. Yellow-highlighted genes were confirmed by qPCR (Fig. 5b). GO term abbreviations are listed in Table in Fig. 5a.

| RefSeq_ID | Nudt1<br>2-KO,<br>FPK<br>M | WT,<br>FPK<br>M | log<sub>2</sub><br>fold<br>change | p value | q value | Gene Symbol | GO<br>Abbrev |
| --- | --- | --- | --- | --- | --- | --- | --- |
| NM_000019 | 79.0 | 29.1 | 1.439 | 5.00E-05 | 2.10E-03 | ACAT1 | Mito |
| NM_005736 | 35.8 | 13.2 | 1.442 | 5.50E-04 | 1.13E-02 | ACTR1A |  |
| NM_000671 | 96.5 | 46.8 | 1.046 | 4.50E-04 | 9.94E-03 | ADH5 | Mito |
| NM_000026 | 147.8 | 65.2 | 1.180 | 1.50E-04 | 4.70E-03 | ADSL | Mito |
| NM_006796 | 39.9 | 13.6 | 1.550 | 1.30E-03 | 1.96E-02 | AFG3L2 | Mito |
| NM_012111 | 128.1 | 62.1 | 1.046 | 7.00E-04 | 1.30E-02 | AHSA1 |  |
| NM_004208 | 46.1 | 21.1 | 1.124 | 3.95E-03 | 4.03E-02 | AIFM1 | Mito |
| NM_016237 | 54.0 | 22.0 | 1.297 | 5.00E-05 | 2.10E-03 | ANAPC5 |  |
| NM_001640 | 47.5 | 23.0 | 1.048 | 1.70E-03 | 2.34E-02 | APEH |  |
| NM_144772 | 150.9 | 59.8 | 1.336 | 1.50E-04 | 4.70E-03 | APOA1BP | Mito |
| NM_004317 | 64.3 | 26.9 | 1.255 | 3.15E-03 | 3.54E-02 | ASNA1 |  |
| NM_018164 | 35.7 | 16.3 | 1.136 | 1.45E-03 | 2.10E-02 | ASUN |  |
| NM_004044 | 78.1 | 26.3 | 1.573 | 5.00E-05 | 2.10E-03 | ATIC | Mito |
| NM_005176 | 255.6 | 106.8 | 1.259 | 3.50E-04 | 8.56E-03 | ATP5G2 |  |
| NM_001697 | 412.9 | 191.1 | 1.111 | 2.00E-04 | 5.73E-03 | ATP5O | Mito |
| NM_170746 | 195.7 | 73.1 | 1.421 | 4.75E-03 | 4.54E-02 | C11orf31 |  |
| NM_005898 | 47.4 | 19.4 | 1.286 | 2.40E-03 | 2.96E-02 | CAPRIN1 |  |
| NM_024098 | 49.4 | 21.3 | 1.212 | 2.85E-03 | 3.30E-02 | CCDC86 | RIB |
| NM_031966 | 100.5 | 37.7 | 1.416 | 5.00E-05 | 2.10E-03 | CCNB1 |  |
| NM_005998 | 291.0 | 140.2 | 1.053 | 1.05E-03 | 1.68E-02 | CCT3 |  |
| NM_006430 | 216.3 | 102.3 | 1.080 | 1.50E-04 | 4.70E-03 | CCT4 |  |
| NM_001255 | 59.8 | 27.8 | 1.106 | 2.55E-03 | 3.08E-02 | CDC20 |  |
| NM_001810 | 37.6 | 17.8 | 1.076 | 1.75E-03 | 2.39E-02 | CENPB |  |
| NM_005441 | 41.4 | 20.1 | 1.042 | 4.55E-03 | 4.42E-02 | CHAF1B |  |
| NM_017801 | 49.3 | 21.0 | 1.235 | 2.00E-04 | 5.73E-03 | CMTM6 |  |
| NM_001206641 | 160.3 | 52.7 | 1.606 | 5.50E-04 | 1.13E-02 | COA6 | Mito |
| NM_004645 | 34.7 | 16.9 | 1.037 | 4.45E-03 | 4.37E-02 | COIL |  |
| NM_006837 | 56.1 | 23.1 | 1.284 | 3.65E-03 | 3.87E-02 | COPS5 |  |
| NM_006833 | 164.7 | 78.8 | 1.063 | 3.00E-04 | 7.72E-03 | COPS6 |  |
| NM_005694 | 336.1 | 128.5 | 1.387 | 1.10E-03 | 1.74E-02 | COX17 | Mito |
| NM_001865 | 474.7 | 234.8 | 1.015 | 2.00E-04 | 5.73E-03 | COX7A2 |  |
| NM_000100 | 81.4 | 21.6 | 1.916 | 2.10E-03 | 2.69E-02 | CSTB |  |
| NM_003798 | 54.3 | 26.3 | 1.046 | 1.50E-03 | 2.15E-02 | CTNNAL1 |  |
| NM_018122 | 63.0 | 23.6 | 1.413 | 1.00E-04 | 3.48E-03 | DARS2 | Mito |
| NM_018959 | 75.4 | 36.2 | 1.059 | 1.00E-03 | 1.62E-02 | DAZAP1 |  |
| NM_017895 | 50.6 | 23.9 | 1.085 | 1.15E-03 | 1.80E-02 | DDX27 | RIB |

| **RefSeq_ID** | **Nudt1**<br>**2-KO,**<br>**FPK**<br>**M** | **WT,**<br>**FPK**<br>**M** | **log<sub>2</sub>**<br>**fold**<br>**change** | **p value** | **q value** | **Gene Symbol** | **GO**<br>**Abbrev** |
| --- | --- | --- | --- | --- | --- | --- | --- |
| NM_006145 | 58.9 | 18.9 | 1.643 | 5.00E-05 | 2.10E-03 | DNAJB1 |  |
| NM_014597 | 51.2 | 23.8 | 1.102 | 1.05E-03 | 1.68E-02 | DNTTIP2 |  |
| NM_005510 | 100.3 | 21.0 | 2.253 | 5.00E-05 | 2.10E-03 | DXO |  |
| NM_006824 | 132.9 | 58.1 | 1.193 | 1.00E-04 | 3.48E-03 | EBNA1BP2 | RIB |
| NM_001398 | 71.9 | 30.4 | 1.243 | 2.55E-03 | 3.08E-02 | ECH1 | Mito |
| NM_004092 | 90.6 | 34.6 | 1.389 | 4.00E-04 | 9.24E-03 | ECHS1 | Mito |
| NM_014740 | 106.4 | 49.1 | 1.116 | 2.00E-04 | 5.73E-03 | EIF4A3 | RIB |
| NM_001418 | 212.6 | 96.4 | 1.142 | 1.95E-03 | 2.54E-02 | EIF4G2 |  |
| NM_006331 | 69.7 | 23.2 | 1.584 | 3.55E-03 | 3.81E-02 | EMG1 | RIB |
| NM_015966 | 122.7 | 53.3 | 1.202 | 6.00E-04 | 1.20E-02 | ERGIC3 |  |
| NM_004730 | 115.8 | 41.7 | 1.473 | 1.00E-04 | 3.48E-03 | ETF1 |  |
| NM_032822 | 87.6 | 37.3 | 1.231 | 4.00E-04 | 9.24E-03 | FAM136A |  |
| NM_002014 | 87.2 | 42.3 | 1.044 | 3.50E-04 | 8.56E-03 | FKBP4 | Mito |
| NM_032020 | 44.3 | 20.6 | 1.105 | 2.35E-03 | 2.93E-02 | FUCA2 |  |
| NM_052850 | 338.3 | 117.8 | 1.522 | 5.00E-05 | 2.10E-03 | GADD45GIP1 | MTE,<br>MTT,<br>Mito |
| NR_002578 | 503.7 | 214.9 | 1.229 | 1.00E-04 | 3.48E-03 | GAS5 |  |
| NM_005271 | 46.2 | 18.2 | 1.340 | 5.00E-05 | 2.10E-03 | GLUD1 | Mito |
| NM_004489 | 74.7 | 30.5 | 1.291 | 2.80E-03 | 3.27E-02 | GPS2 |  |
| NM_000581 | 571.3 | 160.3 | 1.834 | 5.00E-05 | 2.10E-03 | GPX1 | Mito |
| NM_002085 | 223.5 | 98.6 | 1.182 | 3.00E-04 | 7.72E-03 | GPX4 | Mito |
| NM_012203 | 88.3 | 36.9 | 1.257 | 1.35E-03 | 1.99E-02 | GRHPR |  |
| NM_002087 | 53.5 | 25.9 | 1.046 | 1.85E-03 | 2.48E-02 | GRN |  |
| NM_006118 | 206.1 | 97.7 | 1.078 | 2.00E-04 | 5.73E-03 | HAX1 | Mito |
| NM_018072 | 53.0 | 20.1 | 1.396 | 5.00E-05 | 2.10E-03 | HEATR1 | RIB,<br>Mito |
| NM_182922 | 45.7 | 19.2 | 1.248 | 5.50E-04 | 1.13E-02 | HEATR3 |  |
| NM_005340 | 345.0 | 139.6 | 1.305 | 3.50E-04 | 8.56E-03 | HINT1 |  |
| NM_021052 | 267.5 | 107.9 | 1.310 | 5.00E-04 | 1.07E-02 | HIST1H2AE |  |
| NM_003529 | 928.1 | 389.2 | 1.254 | 5.00E-05 | 2.10E-03 | HIST1H3A |  |
| NM_003536 | 2208.1 | 861.1 | 1.359 | 5.00E-05 | 2.10E-03 | HIST1H3H |  |
| NM_175065 | 637.3 | 301.6 | 1.079 | 5.00E-04 | 1.07E-02 | HIST2H2AB |  |
| NM_001123375 | 483.0 | 206.3 | 1.227 | 2.00E-04 | 5.73E-03 | HIST2H3D |  |
| NM_005346_4 | 212.3 | 74.0 | 1.520 | 5.00E-05 | 2.10E-03 | HSPA1B |  |
| NM_002154 | 125.5 | 55.2 | 1.184 | 5.00E-05 | 2.10E-03 | HSPA4 |  |
| NM_016545 | 40.2 | 18.3 | 1.139 | 3.65E-03 | 3.87E-02 | IER5 |  |
| NM_006546 | 54.8 | 22.9 | 1.260 | 6.50E-04 | 1.25E-02 | IGF2BP1 |  |
| NM_006083 | 85.8 | 42.1 | 1.026 | 1.20E-03 | 1.85E-02 | IK |  |
| NM_004515 | 241.8 | 112.8 | 1.100 | 4.00E-04 | 9.24E-03 | ILF2 |  |
| NM_006694 | 59.2 | 18.3 | 1.689 | 3.60E-03 | 3.84E-02 | JTB | Mito |
| NM_017822 | 36.8 | 12.2 | 1.589 | 1.95E-03 | 2.54E-02 | KANSL2 |  |
| NM_015634 | 41.1 | 11.1 | 1.896 | 3.00E-04 | 7.72E-03 | KIAA1279 | Mito |
| NM_015315 | 44.7 | 21.9 | 1.031 | 3.50E-04 | 8.56E-03 | LARP1 |  |
| NM_015340 | 35.3 | 11.6 | 1.609 | 5.05E-03 | 4.73E-02 | LARS2 | Mito |
| NM_018509 | 78.3 | 38.0 | 1.044 | 2.15E-03 | 2.73E-02 | LRRC59 |  |
| NM_152344 | 107.9 | 44.3 | 1.286 | 5.00E-05 | 2.10E-03 | LSM12 |  |
| NM_005911 | 73.8 | 32.8 | 1.171 | 1.25E-03 | 1.90E-02 | MAT2A |  |
| NM_005915 | 59.0 | 28.9 | 1.031 | 5.00E-04 | 1.07E-02 | MCM6 |  |
| NM_014175 | 72.1 | 21.5 | 1.746 | 1.50E-04 | 4.70E-03 | MRPL15 | MTE,<br>MTT,<br>Mito |
| NM_017971 | 200.6 | 84.6 | 1.245 | 2.40E-03 | 2.96E-02 | MRPL20 | MTE,<br>MTT,<br>Mito |
| NM_181514 | 300.3 | 150.0 | 1.002 | 3.80E-03 | 3.94E-02 | MRPL21 | MTE,<br>MTT,<br>Mito |
| NM_016504 | 143.6 | 52.5 | 1.453 | 1.60E-03 | 2.26E-02 | MRPL27 | MTE,<br>MTT,<br>Mito |
| NM_006428 | 111.2 | 46.7 | 1.252 | 5.00E-04 | 1.07E-02 | MRPL28 | MTE,<br>MTT,<br>Mito |
| NM_032479 | 156.3 | 36.9 | 2.084 | 1.20E-03 | 1.85E-02 | MRPL36 | MTE,<br>MTT,<br>Mito |
| NM_022839 | 53.6 | 20.5 | 1.391 | 2.75E-03 | 3.22E-02 | MRPS11 | MTE,<br>MTT,<br>Mito |
| NM_016065 | 89.0 | 41.4 | 1.104 | 9.00E-04 | 1.51E-02 | MRPS16 | MTE,<br>MTT,<br>Mito |
| NM_015969 | 148.0 | 50.7 | 1.546 | 1.95E-03 | 2.54E-02 | MRPS17 | MTE,<br>MTT,<br>Mito |
| NM_016070 | 81.5 | 27.9 | 1.545 | 1.00E-04 | 3.48E-03 | MRPS23 | MTE,<br>MTT,<br>Mito |
| NM_000251 | 42.5 | 21.2 | 1.002 | 2.15E-03 | 2.73E-02 | MSH2 |  |
| NM_002444 | 41.7 | 18.8 | 1.146 | 2.50E-04 | 6.97E-03 | MSN |  |
| NM_004739 | 55.6 | 21.7 | 1.358 | 5.00E-05 | 2.10E-03 | MTA2 |  |
| NM_079423 | 540.9 | 211.8 | 1.353 | 9.00E-04 | 1.51E-02 | MYL6 |  |
| NM_015965 | 390.6 | 194.9 | 1.003 | 7.50E-04 | 1.35E-02 | NDUFA13 | Mito |
| NM_002489 | 46.0 | 16.9 | 1.445 | 1.00E-03 | 1.62E-02 | NDUFA4 | Mito |
| NM_014222 | 228.8 | 92.6 | 1.305 | 5.00E-05 | 2.10E-03 | NDUFA8 | Mito |
| NM_005002 | 79.2 | 38.6 | 1.036 | 2.45E-03 | 3.00E-02 | NDUFA9 | Mito |
| NM_004546 | 255.7 | 107.8 | 1.247 | 3.95E-03 | 4.03E-02 | NDUFB2 | Mito |
| NM_005005 | 151.3 | 59.5 | 1.345 | 5.10E-03 | 4.74E-02 | NDUFB9 | Mito |
| NM_004551 | 133.6 | 41.9 | 1.674 | 1.15E-03 | 1.80E-02 | NDUFS3 | Mito |
| NM_021074 | 100.1 | 45.2 | 1.148 | 3.20E-03 | 3.58E-02 | NDUFV2 | Mito |
| NM_017838 | 278.2 | 107.1 | 1.377 | 5.00E-05 | 2.10E-03 | NHP2 |  |
| NM_032390 | 64.5 | 30.9 | 1.062 | 5.10E-03 | 4.74E-02 | NIFK |  |
| NM_014062 | 68.7 | 29.0 | 1.246 | 4.00E-04 | 9.24E-03 | NOB1 | RIB |
| NM_006310 | 39.7 | 19.1 | 1.061 | 6.00E-04 | 1.20E-02 | NPEPPS |  |
| NM_001134231 | 93.2 | 43.2 | 1.108 | 2.65E-03 | 3.16E-02 | NT5DC2 |  |
| NM_153485 | 41.7 | 20.4 | 1.027 | 7.50E-04 | 1.35E-02 | NUP155 |  |
| NM_024844 | 46.2 | 20.2 | 1.193 | 1.80E-03 | 2.44E-02 | NUP85 |  |
| NM_002533 | 35.6 | 17.4 | 1.030 | 3.40E-03 | 3.72E-02 | NVL | Mito |
| NM_002539 | 169.0 | 50.2 | 1.751 | 1.00E-04 | 3.48E-03 | ODC1 |  |
| NM_032346 | 54.4 | 12.5 | 2.122 | 3.35E-03 | 3.69E-02 | PDCD2L |  |
| NM_020992 | 49.5 | 15.9 | 1.636 | 1.95E-03 | 2.54E-02 | PDLIM1 |  |
| NM_002622 | 46.3 | 13.8 | 1.751 | 5.05E-03 | 4.73E-02 | PFDN1 |  |
| NM_002627 | 45.7 | 20.5 | 1.154 | 1.20E-03 | 1.85E-02 | PFKP |  |
| NM_001144831 | 319.1 | 118.8 | 1.425 | 1.70E-03 | 2.34E-02 | PHB2 | Mito |
| NM_005030 | 114.5 | 51.0 | 1.167 | 5.00E-05 | 2.10E-03 | PLK1 |  |
| NM_000938 | 73.6 | 36.3 | 1.018 | 3.00E-04 | 7.72E-03 | POLR2B |  |
| NM_002696 | 111.4 | 45.0 | 1.307 | 2.70E-03 | 3.19E-02 | POLR2G |  |
| NM_005837 | 86.5 | 35.7 | 1.278 | 4.85E-03 | 4.62E-02 | POP7 |  |
| NM_005729 | 63.9 | 21.3 | 1.586 | 5.00E-05 | 2.10E-03 | PPIF | Mito |
| NM_177983 | 108.6 | 41.4 | 1.392 | 5.00E-05 | 2.10E-03 | PPM1G |  |
| NM_005973 | 55.5 | 17.6 | 1.658 | 1.50E-04 | 4.70E-03 | PRCC |  |
| NM_006406 | 283.5 | 98.7 | 1.522 | 5.00E-05 | 2.10E-03 | PRDX4 | Mito |
| NM_004905 | 112.3 | 54.9 | 1.034 | 7.50E-04 | 1.35E-02 | PRDX6 |  |
| NM_002797 | 124.1 | 53.3 | 1.219 | 1.35E-03 | 1.99E-02 | PSMB5 |  |
| NM_002804 | 92.5 | 41.4 | 1.160 | 5.50E-04 | 1.13E-02 | PSMC3 |  |
| NM_002805 | 146.5 | 55.8 | 1.393 | 2.10E-03 | 2.69E-02 | PSMC5 |  |

| **RefSeq_ID** | **Nudt1**<br>**2-KO,**<br>**FPK**<br>**M** | **WT,**<br>**FPK**<br>**M** | **log<sub>2</sub>**<br>**fold**<br>**change** | **p value** | **q value** | **Gene Symbol** | **GO**<br>**Abbrev** |
| --- | --- | --- | --- | --- | --- | --- | --- |
| NM_002807 | 74.1 | 35.0 | 1.080 | 4.50E-04 | 9.94E-03 | PSMD1 |  |
| NM_002808 | 71.0 | 25.5 | 1.477 | 5.00E-05 | 2.10E-03 | PSMD2 |  |
| NM_006423 | 110.4 | 47.9 | 1.204 | 3.70E-03 | 3.90E-02 | RABAC1 |  |
| NM_018077 | 34.5 | 8.8 | 1.963 | 5.00E-04 | 1.07E-02 | RBM28 |  |
| NM_080748 | 351.0 | 137.2 | 1.355 | 7.00E-04 | 1.30E-02 | ROMO1 | Mito |
| NM_003683 | 52.1 | 22.2 | 1.229 | 2.30E-03 | 2.88E-02 | RRP1 | RIB |
| NM_015169 | 36.4 | 12.5 | 1.538 | 4.75E-03 | 4.54E-02 | RRS1 |  |
| NM_003707 | 128.3 | 60.5 | 1.084 | 3.00E-04 | 7.72E-03 | RUVBL1 |  |
| NM_006666 | 159.8 | 75.2 | 1.087 | 2.00E-04 | 5.73E-03 | RUVBL2 |  |
| NM_138421 | 68.8 | 32.5 | 1.082 | 2.80E-03 | 3.27E-02 | SAAL1 |  |
| NM_015380 | 47.2 | 17.6 | 1.423 | 1.65E-03 | 2.29E-02 | SAMM50 | Mito |
| NM_016038 | 67.9 | 28.7 | 1.242 | 1.35E-03 | 1.99E-02 | SBDS | RIB |
| NR_003023 | 828.1 | 382.3 | 1.115 | 4.50E-04 | 9.94E-03 | SCARNA2 |  |
| NM_004589 | 68.5 | 23.7 | 1.533 | 1.50E-04 | 4.70E-03 | SCO1 | Mito |
| NM_003000 | 119.6 | 58.4 | 1.033 | 1.80E-03 | 2.44E-02 | SDHB | Mito |
| NM_003002 | 117.4 | 56.3 | 1.060 | 1.75E-03 | 2.39E-02 | SDHD | Mito |
| NM_021237 | 105.1 | 17.5 | 2.588 | 9.50E-04 | 1.57E-02 | SELK |  |
| NM_006842 | 102.8 | 44.1 | 1.223 | 5.00E-05 | 2.10E-03 | SF3B2 |  |
| NM_005850 | 46.2 | 19.6 | 1.241 | 2.55E-03 | 3.08E-02 | SF3B4 |  |
| NM_031287 | 160.9 | 76.5 | 1.073 | 3.20E-03 | 3.58E-02 | SF3B5 |  |
| NM_015327 | 39.6 | 18.2 | 1.123 | 1.00E-04 | 3.48E-03 | SMG5 |  |
| NM_020197 | 41.0 | 9.5 | 2.118 | 2.25E-03 | 2.84E-02 | SMYD2 |  |
| NR_003697 | 82.8 | 16.9 | 2.296 | 3.80E-03 | 3.94E-02 | SNHG15 |  |
| NR_132114 | 306.3 | 72.4 | 2.081 | 1.95E-03 | 2.54E-02 | SNHG19 |  |
| NM_014014 | 58.6 | 27.1 | 1.111 | 1.50E-04 | 4.70E-03 | SNRNP200 |  |
| NM_152233 | 42.8 | 21.3 | 1.007 | 1.95E-03 | 2.54E-02 | SNX6 |  |
| NM_003132 | 109.1 | 47.8 | 1.189 | 3.50E-04 | 8.56E-03 | SRM |  |
| NM_003017 | 130.6 | 55.3 | 1.239 | 2.00E-04 | 5.73E-03 | SRSF3 |  |
| NM_020151 | 100.7 | 39.5 | 1.349 | 5.00E-05 | 2.10E-03 | STARD7 | Mito |
| NM_003849 | 51.6 | 21.1 | 1.292 | 4.30E-03 | 4.28E-02 | SUCLG1 | Mito |
| NM_016614 | 40.7 | 15.5 | 1.397 | 2.30E-03 | 2.88E-02 | TDP2 |  |
| NR_001566 | 237.8 | 95.9 | 1.310 | 3.10E-03 | 3.50E-02 | TERC |  |
| NM_018975 | 50.2 | 24.7 | 1.021 | 4.40E-03 | 4.33E-02 | TERF2IP |  |
| NM_016589 | 107.0 | 42.8 | 1.323 | 1.00E-04 | 3.48E-03 | TIMMDC1 | Mito |
| NM_018475 | 37.2 | 14.5 | 1.360 | 4.20E-03 | 4.21E-02 | TMEM165 |  |
| NM_020243 | 125.4 | 57.3 | 1.129 | 4.00E-04 | 9.24E-03 | TOMM22 | Mito |
| NM_006809 | 36.3 | 14.8 | 1.298 | 4.40E-03 | 4.33E-02 | TOMM34 |  |
| NM_007027 | 36.3 | 17.7 | 1.031 | 5.50E-04 | 1.13E-02 | TOPBP1 |  |
| NM_004593 | 81.5 | 31.7 | 1.363 | 5.00E-05 | 2.10E-03 | TRA2B |  |
| NM_001042462 | 102.3 | 33.4 | 1.615 | 4.40E-03 | 4.33E-02 | TRAPPC5 |  |
| NM_006510 | 51.9 | 21.4 | 1.277 | 4.00E-04 | 9.24E-03 | TRIM27 |  |
| NM_015679 | 57.3 | 23.5 | 1.287 | 2.85E-03 | 3.30E-02 | TRUB2 |  |
| NM_018128 | 38.1 | 19.0 | 1.004 | 8.00E-04 | 1.41E-02 | TSR1 | RIB |
| NM_001070 | 79.2 | 26.2 | 1.596 | 2.45E-03 | 3.00E-02 | TUBG1 |  |
| NM_003348 | 69.6 | 24.4 | 1.515 | 5.00E-05 | 2.10E-03 | UBE2N |  |
| NM_004181 | 192.0 | 62.0 | 1.632 | 5.00E-05 | 2.10E-03 | UCHL1 |  |
| NM_003366 | 85.2 | 42.3 | 1.008 | 6.50E-04 | 1.25E-02 | UQCRC2 | Mito |
| NM_014402 | 160.4 | 79.3 | 1.017 | 7.00E-04 | 1.30E-02 | UQCRQ | Mito |
| NM_000375 | 68.6 | 23.6 | 1.542 | 5.00E-04 | 1.07E-02 | UROS | Mito |
| NM_016001 | 62.7 | 29.1 | 1.106 | 2.05E-03 | 2.64E-02 | UTP18 | RIB |
| NM_022916 | 40.1 | 19.5 | 1.037 | 3.85E-03 | 3.98E-02 | VPS33A |  |
| NM_018206 | 58.3 | 23.5 | 1.312 | 5.00E-05 | 2.10E-03 | VPS35 | Mito |
| NM_013245 | 52.1 | 23.8 | 1.133 | 1.65E-03 | 2.29E-02 | VPS4A |  |
| NM_016312 | 64.0 | 31.5 | 1.023 | 1.25E-03 | 1.90E-02 | WBP11 | RIB |
| NM_006784 | 48.9 | 18.7 | 1.386 | 5.00E-05 | 2.10E-03 | WDR3 | RIB |
| NM_032168 | 50.7 | 20.0 | 1.345 | 1.50E-04 | 4.70E-03 | WDR75 | RIB |
| NM_001469 | 241.9 | 117.0 | 1.048 | 3.15E-03 | 3.54E-02 | XRCC6 |  |
| NM_003904 | 70.5 | 25.1 | 1.490 | 2.00E-04 | 5.73E-03 | ZPR1 |  |

**Supplementary Table 3.**
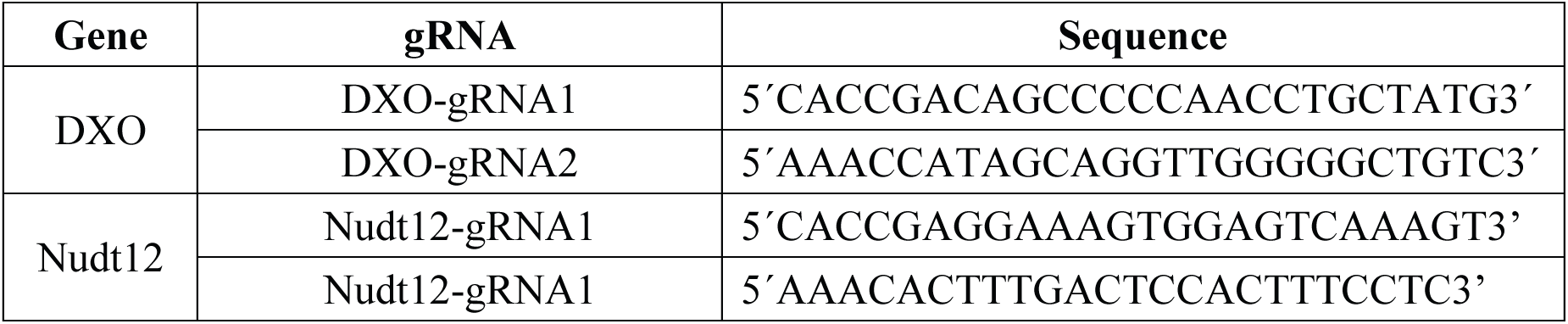

**Supplementary Table 4.**
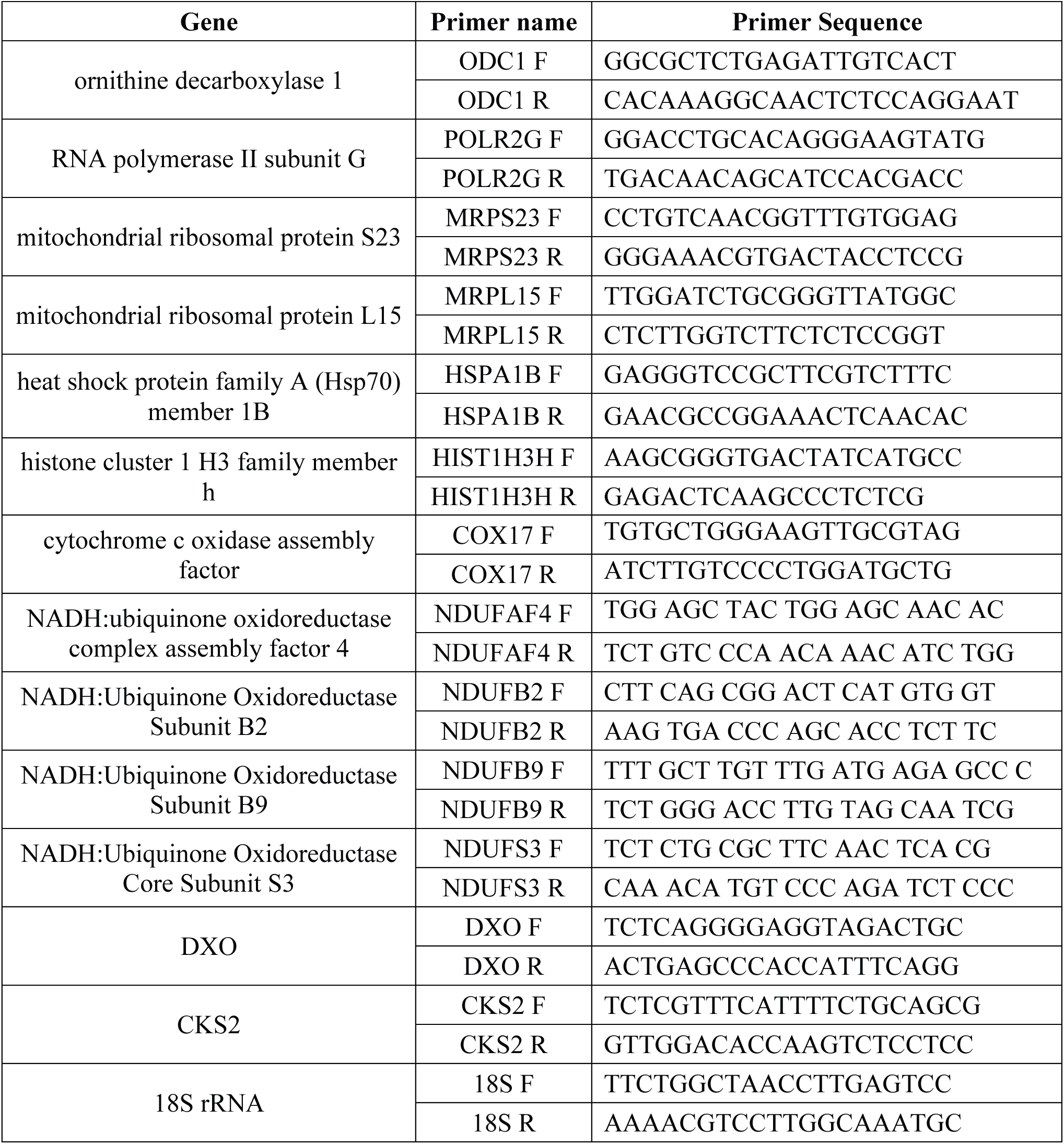
Primers list.

